# Empirical estimation of multiple-testing burden for population-based HLA association studies using sequencing-derived HLA alleles across genetic ancestries

**DOI:** 10.64898/2026.07.12.738059

**Authors:** Daniel Taliun, Sarah A Gagliano Taliun

## Abstract

As population-scale whole-genome sequencing datasets continue to expand, they enable genetic association studies beyond single-nucleotide variants to more complex forms of genetic variation, including classical human leukocyte antigen (HLA) alleles. The HLA region comprises nine highly polymorphic classical HLA genes in extensive linkage disequilibrium that are associated with numerous autoimmune and infectious diseases. However, unlike genome-wide association studies of single-nucleotide variants, there is no general guidance for controlling the multiple-testing burden in HLA allele association analyses. Here, we systematically evaluated the effective number of independent HLA allele tests using sequencing data from diverse genetic ancestries, analytical derivation and simulations. We show that the multiple-testing burden depends on genetic ancestry, allele frequency, and the phenotype model, but remains remarkably stable across minor allele count thresholds, corresponding to approximately 60–70% of the total number of tested HLA alleles. Simulations further demonstrate that the effective number of tests can exceed 90% under realistic disease models. Analyses of 4-field HLA alleles from long-read sequencing showed that higher typing resolution increases the number of alleles but preserves the underlying correlation structure and scales the effective number of independent tests proportionally. Our results provide practical guidance for HLA association studies and support Bonferroni correction based on the total number of tested HLA alleles as a simple and robust approximation when permutation-based approaches are impractical.

---

Genome-wide association studies (GWAS) have successfully identified genetic variants across the genome that are statistically associated with a wide range of human diseases and disease-related traits. Over twenty years ago, three independent research groups deduced that 5x10^-8^, which accounts for 1,000,000 independent tests, was an appropriate statistical significance threshold to account for the multiple testing burden presented by GWAS.^1–3^ Of course, technology, as well as the size and genetic diversity of the available datasets, have advanced considerably since then. One such advancement is significantly improved calling and/or imputing of genetic variation within the Human Leukocyte Antigen (HLA) region on chromosome 6, encoding for a series of immune-related proteins, and notorious for its long-range and complex linkage disequilibrium patterns.

An increasing number of population-scale studies that assess HLA alleles from sequencing data and perform hypothesis-free disease association scans of these alleles, highlight important methodological gaps, including how to account for the multiple-testing burden in this genomic region. Indeed, several biobanks, composed of data from hundreds of thousands of participants each, including FinnGen, UK Biobank and All of Us,^4–7^ have HLA alleles called or imputed, permitting association at scale across a range of phenotypes. However, given the unique correlation patterns within this region, to our knowledge, an investigation of a robust multiple testing threshold has not yet been thoroughly investigated. In this work, we aim to fill this gap by identifying the appropriate HLA-wide significance threshold for classical HLA allele association testing across genetic ancestries. To do so, we systemically assess the number of independent tests in the HLA region through applying complementary approaches at different allele resolutions and different sequencing technologies.

We analyzed 283 2-field alleles in eight of the nine classical Class I and Class II HLA genes across five genetic ancestries in the 1000 Genomes Project Phase 3 high-depth short-read sequencing dataset (**Supplementary Methods**). We restricted analyses to alleles observed at least five times in both the full cohort and in each of the super-populations (minor allele count (MAC) ≥5) (**Table 1**).

**Table 1.** Number of 2-field alleles in classical HLA genes typed in the 1000 Genomes Project using HLA-HD. Alleles presented in this table passed our quality control. MAC – minor allele count. African (AFR, N = 661), Admixed American (AMR, N = 347), East Asian (EAS, N = 504), European (EUR, N = 503), South Asian (SAS, N = 489), and the full cohort (ALL, N = 2,504)

| Gene (class) | Number of alleles with MAC $\geq 1$ | | | | | | Number of alleles with MAC $\geq 5$ | | | | | |
| --- | --- | --- | --- | --- | --- | --- | --- | --- | --- | --- | --- | --- |
|  | AFR | AMR | EAS | EUR | SAS | ALL | AFR | AMR | EAS | EUR | SAS | ALL |
| *A (Class I)* | 44 | 44 | 31 | 29 | 32 | 86 | 26 | 23 | 18 | 18 | 20 | 44 |
| *B (Class I)* | 74 | 98 | 68 | 61 | 58 | 176 | 36 | 38 | 34 | 32 | 32 | 86 |
| *C (Class I)* | 39 | 35 | 32 | 30 | 33 | 74 | 23 | 22 | 19 | 18 | 22 | 35 |
| *DPA1 (Class II)* | 17 | 10 | 9 | 7 | 10 | 27 | 6 | 5 | 5 | 6 | 6 | 12 |
| *DPB1 (Class II)* | 41 | 31 | 31 | 27 | 29 | 74 | 17 | 17 | 15 | 19 | 17 | 36 |
| *DQA1 (Class II)* | -- | -- | -- | -- | -- | -- | -- | -- | -- | -- | -- | -- |
| *DQB1 (Class II)* | 21 | 20 | 17 | 18 | 21 | 36 | 14 | 15 | 15 | 14 | 15 | 18 |
| *DRA (Class II)* | 3 | 3 | 4 | 4 | 3 | 5 | 3 | 2 | 3 | 2 | 3 | 3 |
| *DRB1 (Class II)* | 42 | 45 | 34 | 38 | 42 | 75 | 23 | 31 | 26 | 24 | 23 | 49 |
| **Total** | **281** | **286** | **226** | **214** | **228** | **553** | **148** | **153** | **135** | **133** | **138** | **283** |

Pairwise correlations among HLA alleles demonstrated a structured but sparse correlation pattern, with the majority of allele pairs exhibiting weak correlations. Average correlations were small in magnitude overall but differed between within-gene and between-gene comparisons: the mean pairwise correlation among within-gene allele pairs was −0.0135 (SE = 0.0003), whereas the mean correlation among between-gene allele pairs was 0.0002 (SE = 0.0003) (**Figure 1A**). Although the average between-gene correlation was close to zero, strongly correlated allele pairs were more common among between-gene than within-gene comparisons, with 0.43% and 0.04% of allele pairs, respectively, exhibiting absolute Pearson correlation coefficients >0.3 (**Figure 1B**). These findings are consistent with the underlying correlation structure visualized in pairwise correlation heatmaps (**Supplementary Figure 1**): negative correlations were concentrated among alleles within the same gene, reflecting competition among alternative alleles at that gene, whereas the strongest positive correlations occurred primarily between alleles from different genes, consistent with the extensive haplotypic structure and long-range linkage disequilibrium that characterize the HLA region. We observed consistent trends across all genetic ancestry groups (**Supplementary Figures 2–5**).

**Figure 1.**
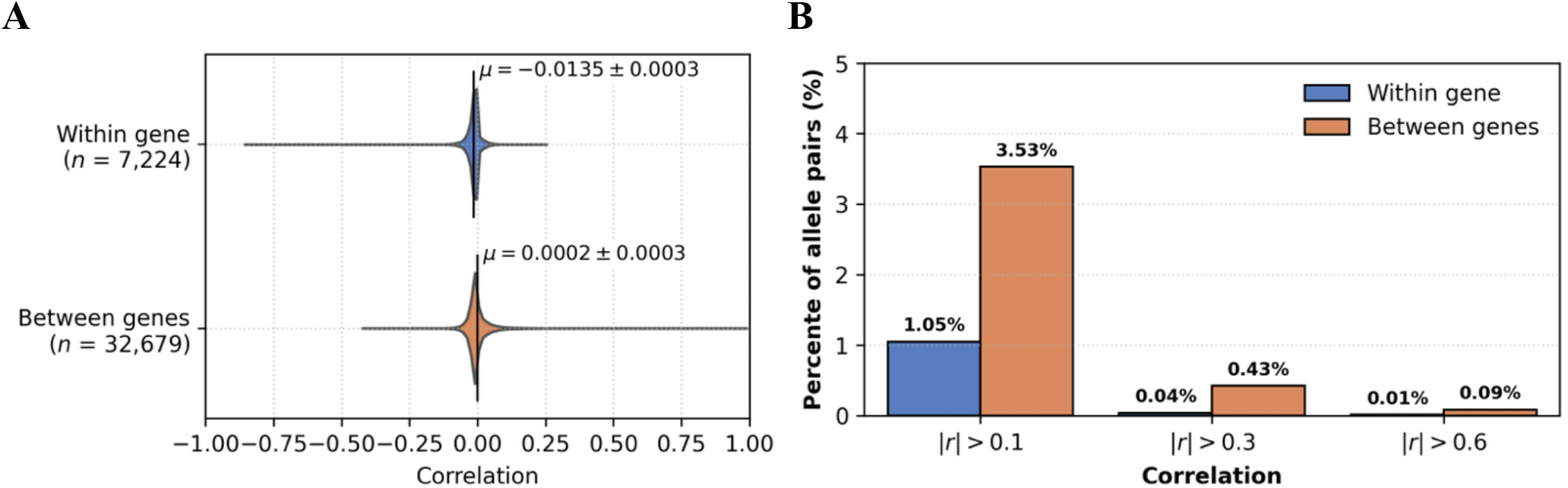
Correlation between 2-field HLA alleles within and between HLA genes. **A**. Distribution of pairwise Pearson correlation coefficients between 2-field HLA alleles within and between HLA genes. Vertical lines indicate the mean correlation coefficient for each distribution, with annotated mean values and standard errors. **B**. Percentage of 2-field HLA allele pairs with pairwise Pearson correlation coefficients exceeding selected thresholds within and between HLA genes. Thresholds were applied to the absolute values of Pearson correlation coefficients (|r|). Bars are ordered by increasing correlation threshold.

**Figure 2.**
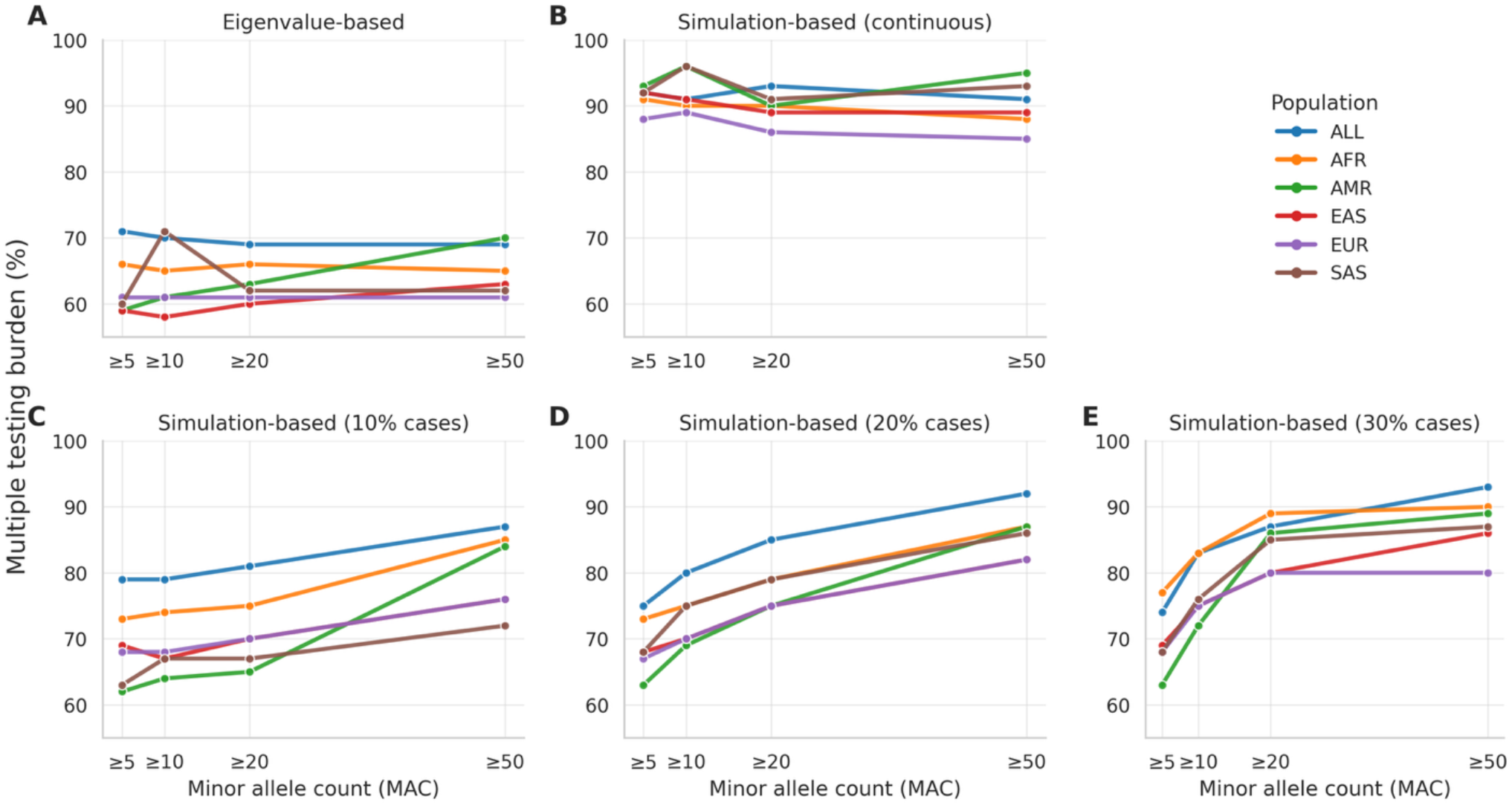
Multiple testing burden for 2-field HLA alleles. Multiple testing burden is defined as *M*_*eff*_/*M*×100%, where *M*_*eff*_ is the effective number of independent tests and *M* is the total number of tested HLA alleles. **A**. *M*_*eff*_ estimated from eigenvalues of the HLA allele correlation matrix. **B**. *M*_*eff*_ estimated from the empirical distribution of minimal p-values obtained from 30,000 simulated continuous phenotypes. **C-E**. *M*_*eff*_ estimated from empirical distributions of minimal p-values obtained from 30,000 simulated binary phenotypes with case prevalences of 10%, 20%, and 30%.

Examples of strongly correlated allele pairs (i.e. haplotypes) across genes included several well-characterized HLA haplotypes (**Supplementary Tables 3-7**). The DRB1*03:01-DQB1*02:01 pair showed high correlation (*r* ≈ 0.96–1.00) across all genetic ancestry groups and has been previously associated with type 1 diabetes risk.^8,9^ Similarly, the C*07:02-B*07:02 pair exhibited strong correlation (*r* ≈ 0.66 in African, *r* ≈ 0.69 Admixed American, and *r* ≈ 0.87 in European genetic ancestry groups) and has been implicated in adverse drug reactions.^10^ Potential population-specific correlations were observed for: DRB1*03:02-DQB1*04:02 pair in African genetic ancestry individuals (*r* ≈ 0.92), previously reported to be protective against type I diabetes in African American individuals;^11^ C*12:02-B*52:01 pair in East Asian genetic ancestry individuals (*r* ≈ 0.78), previously reported to be protective against HIV-1 in treatment-naive Japanese individuals;^12^ and DRB1*08:02-DQB1*04:02 in Admixed American genetic ancestry individuals previously shown to be clinically relevant in studies carried out in Mexico.^13,14^ The differences in correlation patterns between genetic ancestries were beyond these examples: the graph-based analyses where highly correlated allele pairs across genes are connected into disjoint graphs showed that the African genetic ancestry group had smaller disjoint graphs than any other group, reaching a maximum disjoint graph size of only 24 alleles compared to 29 in South Asian, 38 in European, and 55–57 in Admixed American and East Asian groups at |*r*| ≥ 0.3 (**Supplementary Table 8**), reflecting its longer evolutionary history and in line with previously reported shorter haplotype architecture. This pattern of sparse but strong correlations, including ancestry-specific structure, motivates the use of eigenvalue- and permutation-based approaches to estimate the effective number of independent tests (*M*_*eff*_) in association studies, rather than relying on allele counts or assumptions of independence.

Principal component analysis (PCA) of the HLA allele correlation matrices across 1000 Genomes genetic ancestry groups showed that, among >130 HLA alleles analyzed, 48–58 PCs each explained more variance than a single original variable (i.e. HLA allele) (Kaiser criterion; eigenvalue λ > 1). When all genetic ancestry groups were analyzed jointly, the number increased to 116 PCs for the combined correlation structure of 283 HLA alleles (**Supplementary Figures 6-7, Supplementary Table 9**).

Consistently, the eigenvalue-based effective number of independent tests 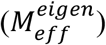, estimated following Li and Ji,^15^ ranged from 79 (East Asian) to 98 (African) across individual ancestries and increased to 202 in the combined dataset. Leave-one-out analyses showed that removal of any single genetic ancestry group resulted in only modest reductions in 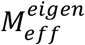(168-190), indicating that no single group disproportionately contributes to the increase in 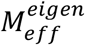. Results were robust across increasingly stringent minor allele count thresholds (MAC ≥5, ≥10 and ≥20), with the combined dataset consistently exhibiting a >2-fold higher 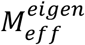than the average across individual groups (**Supplementary Table 9**). Notably, across all genetic ancestry groups and MAC thresholds, 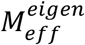remained approximately 59-71% of the total number of alleles (**Figure 2A, Supplementary Table 10**), suggesting a relatively stable degree of correlation-induced redundancy across the analyzed genetic ancestry groups and allele frequency thresholds. Together, these findings indicate that combining genetic ancestries substantially increases the effective dimensionality of HLA allele correlation structure, reflecting correlation patterns that are largely shared but partially distinct across populations.

Next, we investigated whether the effective dimensionality captured by 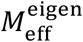translates into the corresponding effective number of independent association tests observed in GWAS under the null hypothesis. We derived empirical simulation-based effective numbers of independent tests from the null distribution of the minimum association p-values across all tested alleles, obtained from 30,000 simulated continuous 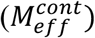 and binary phenotypes 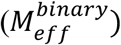. For linear regression, simulation-based estimates were substantially higher than eigenvalue-based estimates, ranging from 117-142 across individual genetic ancestry groups and reaching 260 in the combined dataset (**Supplementary Table 11**). Nevertheless, 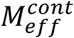consistently represented approximately 88-96% of the total number of tested alleles (**Figure 2B**). In contrast, for binary traits analyzed using Firth logistic regression, estimates at MAC≥5 were more similar, but consistently higher, than the corresponding eigenvalue-based values, ranging from 87 to 108 across ancestry groups and reaching 223 in combined dataset (**Supplementary Table 12**). With increasing MAC thresholds and case prevalence, 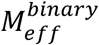increased and approached >90% of the total number of alleles (**Figure 3C-3E**). Although absolute estimates differed between eigenvalue- and simulation-based approaches, the relative inflation in effective testing burden when combining ancestry groups was consistent across methods: all approaches identified a comparable >2-fold increase in effective testing burden when ancestry groups were analyzed jointly.

To assess how HLA typing resolution affects the multiple testing burden, we analyzed 576 distinct 4-field HLA alleles (MAC≥1, 273 with MAC≥5) across nine classical Class I and Class II HLA genes in 415 individuals representing different genetic ancestries from the 1000 Genomes Project long-read sequencing dataset (**Supplementary Methods**). Increasing HLA typing resolution from 2-field to 4-field increased the number of distinct alleles by 2.4-fold for alleles with MAC≥1 and 1.6-fold for alleles with MAC≥5 (**Supplementary Table 13**), indicating that finer-resolution typing primarily partitions common 2-field alleles into multiple less-frequent alleles. The magnitude of this increase varied across genes, with *DRA* exhibiting the largest increase in allele diversity (11-to 14-fold), consistent with its highly conserved coding sequence relative to other classical HLA genes ^16^. The increase in the number of 4-field HLA alleles did not substantially alter the correlation structure observed in the main analyses of 2-field HLA alleles. The strongest correlations remained between alleles from different HLA genes, whereas correlations between alleles within the same gene were generally weaker in absolute magnitude. Overall, most correlations between 4-field HLA alleles were weak, with only ∼1% exhibiting absolute Pearson correlation coefficients >0.3 (**Supplementary Figure 8**). Strongest 4-field HLA allele correlations were consistent with those observed in our main analyses at 2-field resolution. For example, the DRB1*03:01-DQB1*02:01 association observed at 2-field resolution was also observed between the corresponding 4-field alleles DRB1*03:01:01:01-DQB1*02:01:01:01, although with reduced correlation strength (*r* ≈ 0.77 vs *r* ≈ 0.61 at 2- and 4-field resolution, respectively), consistent with partitioning of broad 2-field allele groups (**Supplementary Tables 13 and 14**). Conversely, the C*07:02-B*07:02 association observed at 2-field resolution was refined at 4-field resolution, with the specific allele pair C*07:02:01:03–B*07:02:01:01 showing stronger correlation than the corresponding 2-field pair (*r* ≈ 0.72 vs *r* ≈ 0.49, respectively) (**Supplementary Tables 14 and 15**). Thus, 4-field HLA correlations capture the same known biological linkage patterns observed at 2-field resolution, while the transition to 4-field alleles can either attenuate or strengthen individual correlation coefficients. The absolute number of strongly correlated 4-field alleles between different HLA genes increased compared with 2-field typing, but, relative to the total number of allele pairs, it remained low as in 2-field typing. The similarities in correlation structure were also reflected in the eigenvalue-based effective number of independent tests 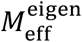. At MAC≥5, it increased from 105 for 2-field HLA alleles to 150 for 4-field HLA alleles, reflecting the larger number of alleles tested (**Supplementary Table 16**). Consistent with our main analyses of 2-field HLA alleles, however, the effective number of independent tests 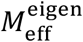remained stable relative to the total number of alleles, ranging from 55 to 68% of the total number of alleles across all MAC thresholds and both typing resolutions. Similarly, the simulation-based effective number of independent tests 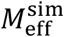, derived from 30,000 simulated continuous phenotypes, increased for 4-field HLA alleles compared to 2-field HLA alleles but remained close to 90% of the total number of tested alleles for both resolutions (**Supplementary Table 17**).

Our estimation of the multiple testing burden using 1000 permutations in the UK Biobank^4^ for two continuous traits ranged from 69%-81% of the total number of tested HLA alleles with MAC > 5, which aligns with our simulation-based results in the 1000 Genomes Project. In particular, using UK Biobank HLA imputation data, we observed 268 independent tests out of 332 tested alleles (i.e. 81% multiple testing burden) for height across 449,744 unrelated participants with non-missing phenotype values. For triglycerides, we observed 229 independent tests out of 332 tested alleles (69%) using 421,488 unrelated participants with non-missing phenotype values.

We found that the multiple-testing burden for HLA association studies depends on allele frequency, genetic ancestry, and phenotype model. Consistent with observations in single nucleotide variant (SNV)-based GWAS, the multiple-testing burden is the highest in the African genetic ancestry group (as defined here as the 1000 Genomes African super-population), which is driven by this group’s longer evolutionary history and shorter haplotype architecture. A critical insight from our work is that pooled analysis of genetic ancestries not only increases the number of tested alleles but also approximately doubles the multiple-testing burden compared to any single ancestry analysis. Consequently, multi-ancestry HLA association studies must account for this increased dimensionality to avoid inflated Type I error rates.

We observed a discrepancy between eigenvalue-based and simulation-based estimates of the effective number of tests. Eigenvalue-based methods suggested a reduction in burden to approximately 60-70% of the total allele count, whereas simulation-based methods, which estimate the effective number of tests empirically from the null distribution, reached >90%. The discrepancy was less pronounced for lower-frequency alleles analyzed in low-prevalence binary phenotypes, where limited statistical information reduces the contribution of rarer alleles to the overall testing burden. This pattern was not observed for continuous traits. More generally, the gap between simulation- and eigenvalue-based estimates likely reflects the sparse correlation structure of the HLA alleles: the vast majority of allele pairs are only weakly correlated and simulation-based methods recognize these as nearly independent tests, whereas eigenvalue-based methods may over-attribute variance to a smaller number of principal components (that is, independent tests).

Our analysis of 4-field HLA alleles derived from long-read sequencing suggests that, as advances in sequencing technologies enable increasingly higher-resolution HLA typing and the discovery of additional rarer alleles, the overall properties of the underlying correlation structure are likely to remain stable. Specifically, although the effective number of independent tests is expected to increase with the total number of alleles tested, its proportion relative to the total number of alleles is expected to remain broadly constant and similar to that observed for 2-field HLA alleles.

In practice, our results suggest that despite extensive linkage disequilibrium in the HLA region, the effective multiple-testing burden in HLA association studies approaches a full Bonferroni correction (i.e., the correction factor is the total number of tested HLA alleles). Across all ancestries and frequency thresholds tested, the effective number of tests for continuous traits corresponded to 88%-96% of the total number of tested alleles, with binary traits approaching similar percentages as allele frequency and case prevalence increased. Therefore, while permutation is always preferred for precision when computationally feasible, we recommend using the total number of tested HLA alleles as a reasonable proxy for establishing significance thresholds in most population-based HLA association analyses.

Our study has several limitations. Despite substantial methodological advances, HLA typing from short-read sequencing data at 2-field resolution remains less accurate than SNV genotyping. Similarly, although long-read sequencing enables 4-field HLA typing, allele calls are not error-free, particularly when generated from earlier ONT flow cells, as used here. Consequently, genotyping errors may introduce some noise into the estimated HLA allele correlation matrices, although we do not expect such errors to alter our main conclusions. At the same time, short-read sequencing remains the predominant technology in large-scale human genetic studies, making our primary analyses representative of the settings in which HLA alleles are used in population-scale association studies. Our analyses also focused on the most commonly used representation of HLA alleles, in which each HLA allele is modelled as a dosage variable. The correlation structure and effective number of independent tests may differ for alternative representations. Furthermore, our simulations used unrelated individuals and correctly specified association models. In practice, violations of these assumptions may inflate type I error even after appropriate multiple-testing correction, underscoring the importance of rigorous quality control and appropriate statistical modelling in HLA studies, as for any association study.

Finally, our work suggests that HLA association studies are unlikely to converge on a universal, technology-independent threshold analogous to the 5x10^-8^ commonly used threshold for SNV-based GWAS. Instead, rather than a fixed constant, the “HLA-wide significance” threshold must be viewed as a dynamic value defined by the specific allelic diversity and ancestral composition of the study cohort.

## Data and code availability

All data used are publicly available. Code and summary-level data are available on GitHub: https://github.com/dtaliun/Multiple_testing_burden_HLA.

## Acknowledgements

This Project has been made possible by the Canada Brain Research Fund (CBRF), an innovative arrangement between the Government of Canada (through Health Canada) and Brain Canada (BCFA_2025_00912), and by the Krembil Foundation. The views expressed herein do not necessarily represent the views of the Minister of Health or the Government of Canada. DT and SAGT also acknowledge the support of the Natural Sciences and Engineering Research Council of Canada (NSERC) Discovery Grant Program: RGPIN-2025-06572 (DT) and RGPIN-2026-04385 (SAGT). SAGT acknowledges salary support from a Fonds de Recherche du Québec – Santé (FRQS; https://frq.gouv.qc.ca) Junior 2 Award (https://doi.org/10.69777/347366). This research was enabled in part by support provided by Calcul Quebec (https://www.calculquebec.ca), the Digital Research Alliance of Canada (https://www.alliancecan.ca). This research has been conducted using the UK Biobank Resource under Application number 66222 (SAGT).

