## Supplementary Information for "Empirical estimation of multiple-testing burden for population-based HLA association studies using sequencing-derived HLA alleles across genetic ancestries"

Table of Contents

**1. Supplementary Methods**
**2. Supplementary References**
**3. Supplementary Tables**
**4. Supplementary Figures**

#### 1. Supplementary Methods

##### *1.1. Sequencing data*

We used 2,504 samples from the 1000 Genomes Project, sequenced by the New York Genome Center using paired-end short-read technology on an Illumina NovaSeq 6000, with an average depth of coverage of 30×. The genetic data were aligned to the GRCh38 human reference genome with HLA decoys and distributed as CRAM files. These CRAM files contained read pairs in which both, one, or neither read was mapped. To improve processing efficiency, we extracted all read pairs with at least one read mapped to the extended HLA region (chr6:28,510,120-33,480,577 or any HLA decoy sequence), as well as read pairs in which neither read was mapped, into smaller intermediate BAM files for subsequent HLA typing.

##### *1.2. 2-field resolution HLA typing*

For HLA typing, we used HLA-HD v1.7.1 with known HLA allele sequences pulled from the IPD-IMGT/HLA database in June 2026. We converted input BAM files to FASTQ files using GATK Picard SamToFastq v4.6.2.0, following the instructions in the HLA-HD documentation. Specifically, we followed the recommended HLA-HD settings: a minimum read length of 100 (-m option) and a trimming rate of 0.95 (-c option). Alleles in the following 29 HLA genes were typed to up to 6 digits: *A*, *B*, *C*, *DRB1*, *DQA1*, *DQB1*, *DPA1*, *DPB1*, *DMA*, *DMB*, *DOA*, *DOB*, *DRA*, *DRB2*, *DRB3*, *DRB4*, *DRB5*, *DRB6*, *DRB7*, *DRB8*, *DRB9*, *E*, *F*, *G*, *H*, *J*, *K*, *L* and *V*.

##### *1.3. 2-field resolution HLA typing quality control*

Quality control of HLA typing was performed separately within each 1000 Genomes genetic ancestry super-population: African (AFR, N = 661), Admixed American (AMR, N = 347), East Asian (EAS, N = 504), European (EUR, N = 503), South Asian (SAS, N = 489), and also in the combined dataset (N = 2,504). First, all typed 3-field HLA alleles were converted to 2-field (i.e. 4-digit) resolution, and any suffixes (e.g., N, L, S, C, Q, A, P, and G) were removed for simplicity. For each sample, an allele pair was considered to have sufficient coverage if both alleles had a minimum coverage >10 and all exons of the corresponding HLA gene were covered. Allele pairs with insufficient coverage were set to missing. HLA genes for which >1% of allele pairs had insufficient coverage were subsequently excluded.

For each sample, an allele pair was considered ambiguous if the coverage ratio between the two alleles was <0.5 (i.e., one allele had more than twice the coverage of the other) or if HLA-HD reported multiple likely candidate allele pairs. Ambiguous allele pairs were set to missing. HLA genes for which >1% of allele pairs were ambiguous were excluded. Monomorphic typed HLA alleles or any that became monomorphic after quality control were excluded.

In total, 627 (329 with minor allele count [MAC] ≥5) alleles across 17 HLA genes consistently passed our quality-control criteria (**Supplementary Tables 1-2**). Of these, 553 (88%) (283 [86%] with MAC≥5) were in eight classical HLA genes: *A*, *B*, *C*, *DPA1, DPB1, DQB1, DRA*, and *DRB1*.

##### *1.4. Correlation, graph-based and principal component analyses*

HLA alleles were encoded in a binary dosage matrix $G \in \mathbb{R}^{n \times a}$, where $n$ denotes the number of samples and $a$ the number of alleles. Matrix entries represented allele dosage per individual (0, 1, or 2 copies; corresponding to genotypes 0/0, 0/1, and 1/1). The pairwise allele correlation matrix $R \in \mathbb{R}^{a \times a}$ was computed from $G$, with elements $r_{ij}$ defined as the Pearson correlation between alleles $i$ and $j$. Hierarchical clustering was performed on $R$ using the seaborn Python package.[1] Graphs of correlated alleles were constructed by treating alleles as nodes and connecting pairs $i, j$ if $|r_{ij}|$ exceeded a predefined threshold. Connected component analysis was then performed using the NetworkX Python package to identify maximal disjoint subgraphs.[2] Principal component analysis (PCA) was applied to $R$, and eigenvalues were computed using NumPy.[3]

*1.5. Population stratification*

We computed the top 10 principal components (PCs) from the variance-standardized genetic relationship matrix for each 1000 Genomes super-population separately, as well as for the combined dataset, using PLINK 2.[4] Only bi-allelic single-nucleotide polymorphisms (SNPs) with a minor allele frequency (MAF) ≥5% were included. Variants were pruned for linkage disequilibrium (LD) using PLINK 2 with a 1,000-kb sliding window, a step size of 100 variants, and an LD threshold of $r^2 > 0.6$. SNPs located within previously defined long-range LD regions,[5] including the HLA region on chromosome 6, were excluded from the PCA. The resulting top 10 PCs were used for phenotype simulations and HLA allele association analyses.

*1.6. Continuous phenotype simulations*

Continuous phenotypes were simulated under a linear model with additive covariate effects. For each individual $i$, the phenotype was generated as

$$y_i = \mu_i + \varepsilon_i$$

where $\varepsilon_i \sim \mathcal{N}(0, 1)$ represents environmental noise. In simulations including covariate effects, the expected phenotype was modeled as

$$\mu_i = 0.1 \, \mathrm{sex}_i + \sum_{k=1}^{10} \beta_k \, \mathrm{PC}_{ik}$$

where sex was coded as a binary covariate and the first 10 PCs captured genetic ancestry. Prior to simulation, all PCs were standardized to zero mean and unit variance. PC effect sizes $\beta_k$ were assigned as a geometrically decreasing sequence ranging from 0.3 for PC1 to 0.01 for PC10, thereby modeling progressively weaker ancestry effects across successive PCs. When covariates were omitted, $\mu_i = 0$ for all individuals. Independent phenotypes were generated by drawing residual errors from a standard normal distribution (i.e. $\sigma = 1$). 30,000 phenotypes were simulated independently using the same covariate effects but different realizations of the residual error.

*1.7. Binary phenotype simulation*

Binary phenotypes were simulated under a logistic regression model. For each individual $i$, disease status was generated as

$$y_i \sim \mathrm{Bernoulli}(p_i)$$

where

$$\mathrm{logit}(p_i) = \beta_0 + \ln(1.5) \, sex_i + \sum_{k=1}^{10} \beta_k \, \mathrm{PC}_{ik}$$

Sex was included as a binary covariate, and the first 10 PCs capturing genetic ancestry as continuous covariates. Prior to simulation, all PCs were standardized to zero mean and unit variance. PC effect sizes were specified as odds ratios forming a geometrically decreasing sequence from 1.4 for PC1 to 1.01 for PC10, with regression coefficients defined as $\beta_k = ln(\mathrm{OR}_k)$. The intercept $\beta_0$ was estimated numerically to ensure that the mean predicted disease probability across all individuals matched the specified target case fraction. Individual disease status was then sampled independently from a Bernoulli distribution with probability $p_i$. In simulations without covariate effects, all individuals were assigned the same disease probability: equal to the target case fraction. 30,000 phenotypes were simulated independently using the same logistic model and different realizations of the Bernoulli sampling process.

*1.8. Association tests*

Association analyses between HLA alleles and simulated continuous phenotypes were performed in R (version 4.5.0) using additive linear regression as implemented in the lm function. HLA alleles were coded as allele dosages (0, 1, or 2 copies), and the following model was fitted separately for each allele-phenotype pair:

$$y_i = \beta_0 + \beta_a G_{ia} + \beta_{sex}\, \mathrm{sex}_i + \sum_{k=1}^{10} \beta_k\, \mathrm{PC}_{ik}$$

where $y_i$ is the phenotype of individual $i$, $G_{ia}$ is the HLA allele $a$ dosage in individual $i$, and $\mathrm{PC}_{ik}$ denotes the first 10 PCs capturing genetic ancestry in individual $i$. Sex was included as a binary variable. All PCs were standardized to zero mean and unit variance prior to analysis. Resulting standard linear regression p-values were used for all downstream analyses.

Association analyses between HLA alleles and simulated binary phenotypes were performed in R (version 4.5.0) using Firth's bias-reduced logistic regression as implemented in the R package logistf.[6] For each allele-phenotype pair, the following model was fitted:

$$\mathrm{logit}(\mathrm{P}(y_i = 1)) = \beta_0 + \beta_a G_{ia} + \beta_{sex}\, \mathrm{sex}_i + \sum_{k=1}^{10} \beta_k\, \mathrm{PC}_{ik}$$

where $y_i$ denotes binary phenotype status for individual $i$, $G_{ia}$ is the HLA allele $a$ dosage in individual $i$, and $\mathrm{PC}_{ik}$ represents the first 10 PCs of genetic ancestry in individual $i$. Sex was included as a binary variable. PCs were standardized to zero mean and unit variance prior to analysis. Binary phenotypes were coded as 0/1 prior to model fitting. Firth's penalized likelihood approach was used to reduce small-sample bias and to improve robustness in the presence of rare variants and case–control imbalance. Resulting penalized likelihood p-values were carried forward for all downstream analyses due to their robustness in the presence of rare alleles and case–control imbalance.

*1.9. Simulation-based estimation of the effective number of independent tests*

To estimate the multiple testing burden under the null, we generated an empirical null distribution of association statistics by simulating 30,000 continuous and 30,000 binary phenotypes under the corresponding null models, preserving genotype structure, followed by association testing across

HLA alleles. For each simulation, the minimum (i.e. most significant) p-value across all tested alleles was recorded. The resulting distribution of minimum p-values was used to estimate its 5th percentile $q_{0.05}(\min p)$. An effective multiple testing correction factor was then defined as $0.05/q_{0.05}(\min p)$.

*2.0. Analyses of 4-field HLA alleles*

We used 4-field HLA alleles typed with SpecImmune[7] from 1,019 individuals in the 1000 Genomes Project sequenced using Oxford Nanopore Technologies (ONT) long-read sequencing (R9.4.1 flow cells). This dataset, comprising individuals of diverse genetic ancestries, has been described previously,[8] and the SpecImmune HLA typing has been described and benchmarked elsewhere. We restricted the dataset to 908 unrelated individuals (as defined by 1000 Genomes) and excluded samples with >5% missing HLA allele calls across the nine classical HLA genes, leaving 415 individuals for analysis. After quality control, only *DPA1* (0.48% missing alleles) and *DRA* (14.22%) retained missing allele calls among the classical HLA genes. *DRA* was retained for completeness because sensitivity analyses showed that its inclusion had negligible effects on the results. HLA allele calls supported by fewer than 10 sequencing reads were set to missing. For comparison with conventional HLA resolution, 2-field alleles were generated by collapsing 4-field alleles sharing the same first two fields. All downstream analyses were performed using the same analytical framework as performed in the main analyses.

*2.1. Permutations of continuous phenotypes in the UK Biobank for imputed HLA alleles*

We then assessed the number of effective tests using permutations of rank-based inverse transformed continuous phenotypes from the UK Biobank measured at the baseline visit: standing height (field 50) and triglyceride levels (field 30870). We used the 2-field HLA best-guess derived genotypes (field 22182) that are available for 11 HLA genes: *A*, *B*, *C, DRB5*, *DRB4*, *DRB3*, *DRB1*, *DQB1*, *DQA1*, *DPB1*, *DPA1*, as previously described.[9,10] Out of the 362 alleles available, we removed those within non-classical HLA genes (*DRB3*, *DRB4*, *DRB5*), which left 349 alleles. We then restricted the dataset to the subset of unrelated individuals (kinship coefficient less than 0.084) and removed HLA alleles with a MAC greater than 5, which left 332 alleles across 488,251 individuals for analysis. We used the commonly used PLINK2 software to carry out association testing for 1000 permutations using standardized covariates (age, sex and the first twenty PCs).

### 3. Supplementary Tables

**Supplementary Table 1. Number of 2-field HLA alleles typed in the 1000 Genomes Project using HLA-HD before quality checks.** African (AFR, N = 661), Admixed American (AMR, N = 347), East Asian (EAS, N = 504), European (EUR, N = 503), South Asian (SAS, N = 489), and the full cohort (ALL, N = 2,504)

| **Gene** | **Number of alleles** | | | | | |
|---|---|---|---|---|---|---|
| | **AFR** | **AMR** | **EAS** | **EUR** | **SAS** | **ALL** |
| *A* | 44 | 44 | 31 | 29 | 32 | 86 |
| *B* | 74 | 98 | 68 | 61 | 58 | 176 |
| *C* | 39 | 35 | 32 | 30 | 33 | 74 |
| *DMA* | 5 | 5 | 4 | 4 | 4 | 5 |
| *DMB* | 6 | 6 | 6 | 6 | 4 | 7 |
| *DOA* | 2 | 5 | 5 | 6 | 5 | 10 |
| *DOB* | 6 | 6 | 5 | 6 | 5 | 7 |
| *DPA1* | 17 | 10 | 9 | 7 | 10 | 27 |
| *DPB1* | 41 | 31 | 31 | 28 | 29 | 75 |
| *DQA1* | 18 | 17 | 18 | 16 | 15 | 26 |
| *DQB1* | 21 | 20 | 17 | 18 | 21 | 36 |
| *DRA* | 3 | 3 | 4 | 4 | 3 | 5 |
| *DRB1* | 42 | 46 | 34 | 38 | 42 | 75 |
| *DRB2* | 1 | 1 | 1 | 1 | 1 | 1 |
| *DRB3* | 8 | 7 | 9 | 8 | 7 | 19 |
| *DRB4* | 4 | 4 | 5 | 3 | 5 | 8 |
| *DRB5* | 4 | 5 | 6 | 4 | 6 | 13 |
| *DRB6* | 3 | 3 | 3 | 3 | 3 | 3 |
| *DRB7* | 1 | 1 | 1 | 1 | 1 | 1 |
| *DRB8* | 1 | 1 | 1 | 1 | 1 | 1 |
| *DRB9* | 2 | 2 | 2 | 2 | 2 | 2 |
| *E* | 9 | 4 | 6 | 7 | 6 | 17 |
| *F* | 4 | 3 | 5 | 4 | 4 | 10 |
| *G* | 9 | 7 | 10 | 6 | 6 | 16 |
| *H* | 19 | 15 | 13 | 16 | 15 | 26 |
| *J* | 1 | 1 | 2 | 1 | 1 | 2 |
| *K* | 4 | 4 | 4 | 4 | 4 | 4 |
| *L* | 3 | 3 | 3 | 3 | 3 | 3 |
| *V* | 1 | 1 | 1 | 1 | 1 | 1 |
| **Total** | **392** | **388** | **336** | **318** | **327** | **736** |

**Supplementary Table 2. Number of 2-field HLA alleles typed in the 1000 Genomes Project using HLA-HD.** Alleles presented in this table passed our quality control. MAC – minor allele count. African (AFR, N = 661), Admixed American (AMR, N = 347), East Asian (EAS, N = 504), European (EUR, N = 503), South Asian (SAS, N = 489), and the full cohort (ALL, N = 2504)

| **Gene (class)** | **Number of alleles with MAC ≥1** | | | | | | **Number of alleles with MAC ≥5** | | | | | |
|---|---|---|---|---|---|---|---|---|---|---|---|---|
| | **AFR** | **AMR** | **EAS** | **EUR** | **SAS** | **ALL** | **AFR** | **AMR** | **EAS** | **EUR** | **SAS** | **ALL** |
| *A (classical Class I)* | 44 | 44 | 31 | 29 | 32 | 86 | 26 | 23 | 18 | 18 | 20 | 44 |
| *B (classical Class I)* | 74 | 98 | 68 | 61 | 58 | 176 | 36 | 38 | 34 | 32 | 32 | 86 |
| *C (classical Class I)* | 39 | 35 | 32 | 30 | 33 | 74 | 23 | 22 | 19 | 18 | 22 | 35 |
| *DMA (Class II)* | 5 | 5 | -- | 4 | 4 | 5 | 3 | 4 | -- | 4 | 4 | 5 |
| *DMB (Class II)* | 6 | 6 | 6 | 6 | 4 | 7 | 5 | 5 | 5 | 5 | 4 | 6 |
| *DOA (Class II)* | 1 | 5 | 5 | 6 | 5 | 9 | 1 | 3 | 2 | 3 | 4 | 5 |
| *DOB (Class II)* | 6 | 6 | 5 | 6 | 5 | 7 | 6 | 5 | 4 | 5 | 4 | 6 |
| *DPA1 (classical Class II)* | 17 | 10 | 9 | 7 | 10 | 27 | 6 | 5 | 5 | 6 | 6 | 12 |
| *DPB1(classical Class II)* | 41 | 31 | 31 | 27 | 29 | 74 | 17 | 17 | 15 | 19 | 17 | 36 |
| *DQA1 (classical Class II)* | -- | -- | -- | -- | -- | -- | -- | -- | -- | -- | -- | -- |
| *DQB1 (classical Class II)* | 21 | 20 | 17 | 18 | 21 | 36 | 14 | 15 | 15 | 14 | 15 | 18 |
| *DRA (classical Class II)* | 3 | 3 | 4 | 4 | 3 | 5 | 3 | 2 | 3 | 2 | 3 | 3 |
| *DRB1 (classical Class II)* | 42 | 45 | 34 | 38 | 42 | 75 | 23 | 31 | 26 | 24 | 23 | 49 |
| *DRB2 (pseudogene Class II)* | -- | -- | -- | -- | -- | -- | -- | -- | -- | -- | -- | -- |
| *DRB3 (secondary classical Class II)* | -- | -- | -- | -- | -- | -- | -- | -- | -- | -- | -- | -- |
| *DRB4 (secondary classical Class II)* | -- | -- | -- | -- | -- | -- | -- | -- | -- | -- | -- | -- |
| *DRB5 (secondary classical Class II)* | -- | -- | -- | -- | -- | -- | -- | -- | -- | -- | -- | -- |
| *DRB6 (pseudogene Class II)* | -- | -- | -- | -- | -- | -- | -- | -- | -- | -- | -- | -- |
| *DRB7 (pseudogene Class II)* | -- | -- | -- | -- | -- | -- | -- | -- | -- | -- | -- | -- |
| *DRB8 (pseudogene Class II)* | -- | -- | -- | -- | -- | -- | -- | -- | -- | -- | -- | -- |
| *DRB9 (pseudogene Class II)* | -- | -- | -- | -- | -- | -- | -- | -- | -- | -- | -- | -- |
| *E (Class I)* | 9 | 4 | 6 | 7 | 4 | 15 | 4 | 2 | 3 | 3 | 2 | 7 |
| *F (Class I)* | 4 | 3 | 5 | 4 | 4 | 10 | 3 | 2 | 4 | 3 | 2 | 6 |
| *G (Class I)* | 9 | 7 | 10 | 6 | 6 | 16 | 6 | 5 | 5 | 5 | 5 | 8 |
| *H (pseudogene Class I)* | -- | -- | -- | 16 | -- | -- | -- | -- | -- | 12 | -- | -- |
| *J (pseudogene Class I)* | -- | -- | 2 | -- | -- | 2 | -- | -- | 0 | -- | -- | -- |
| *K (pseudogene Class I)* | -- | -- | -- | -- | -- | -- | -- | -- | -- | -- | -- | -- |
| *L (pseudogene Class I)* | 3 | 3 | 3 | 3 | -- | 3 | 3 | 3 | 3 | 3 | -- | 3 |
| *V (pseudogene Class I)* | -- | -- | -- | -- | -- | -- | -- | -- | -- | -- | -- | -- |
| **Total** | **324** | **325** | **268** | **272** | **260** | **627** | **179** | **182** | **161** | **176** | **163** | **329** |

**Supplementary Table 3. Top positively correlated 2-field HLA allele pairs in the African genetic ancestry group (Pearson *r* > 0.6).**

| **Allele 1** | **Allele 2** | **Allele 1 count** | **Allele 2 count** | **Pearson *r*** |
|---|---|---|---|---|
| DPA1*02:12 | DPB1*85:01 | 24 | 26 | 0.96 |
| DRB1*03:01 | DQB1*02:01 | 63 | 64 | 0.96 |
| DRB1*03:02 | DQB1*04:02 | 101 | 102 | 0.93 |
| DRA*01:03 | DRB1*03:02 | 97 | 101 | 0.93 |
| DPA1*03:05 | DPB1*40:01 | 24 | 20 | 0.92 |
| C*02:10 | B*15:03 | 89 | 82 | 0.91 |
| DRA*01:03 | DQB1*04:02 | 97 | 102 | 0.90 |
| DPA1*03:01 | DPB1*105:01 | 167 | 151 | 0.87 |
| C*14:02 | B*15:16 | 14 | 16 | 0.87 |
| C*17:01 | B*42:01 | 96 | 67 | 0.83 |
| C*18:01 | B*81:01 | 14 | 28 | 0.75 |
| DRB1*15:03 | DQB1*06:02 | 156 | 237 | 0.74 |
| C*12:03 | B*39:10 | 16 | 12 | 0.72 |
| DRB1*14:54 | DQB1*05:03 | 18 | 14 | 0.69 |
| C*07:01 | B*49:01 | 82 | 39 | 0.68 |
| C*07:02 | B*07:02 | 65 | 73 | 0.66 |
| DRB1*16:02 | DQB1*05:02 | 20 | 46 | 0.65 |
| DPA1*01:03 | DPB1*02:01 | 346 | 169 | 0.63 |
| DRB1*13:02 | DQB1*06:09 | 103 | 57 | 0.63 |
| DPA1*02:02 | DPB1*01:01 | 238 | 455 | 0.63 |
| DRB1*04:05 | DQB1*03:02 | 23 | 47 | 0.61 |
| DRB1*07:01 | DQB1*02:02 | 87 | 195 | 0.61 |
| C*06:02 | B*58:02 | 95 | 39 | 0.61 |

**Supplementary Table 4. Top positively correlated 2-field HLA allele pairs in the Admixed American genetic ancestry group (Pearson *r* > 0.6).**

| Allele 1 | Allele 2 | Allele 1 count | Allele 2 count | Pearson *r* |
|---|---|---|---|---|
| DRB1*15:02 | DQB1*06:01 | 8 | 8 | 1.00 |
| C*02:10 | B*15:03 | 5 | 5 | 1.00 |
| DRB1*03:01 | DQB1*02:01 | 44 | 46 | 0.98 |
| DRB1*13:01 | DQB1*06:03 | 35 | 34 | 0.95 |
| C*08:02 | B*14:02 | 38 | 36 | 0.91 |
| C*07:18 | B*58:01 | 6 | 5 | 0.91 |
| DPA1*02:07 | DPB1*19:01 | 9 | 8 | 0.88 |
| DRB1*07:01 | DQB1*02:02 | 71 | 68 | 0.87 |
| DPA1*03:01 | DPB1*105:01 | 13 | 15 | 0.87 |
| DRB1*15:01 | DQB1*06:02 | 32 | 40 | 0.82 |
| DRB1*09:01 | DQB1*03:03 | 42 | 47 | 0.79 |
| DRB1*08:02 | DQB1*04:02 | 47 | 74 | 0.79 |
| DRB1*11:02 | DQB1*03:19 | 8 | 6 | 0.77 |
| C*14:02 | B*15:16 | 7 | 6 | 0.77 |
| DRB1*13:02 | DQB1*06:04 | 14 | 11 | 0.75 |
| C*08:01 | B*48:01 | 20 | 21 | 0.73 |
| C*12:02 | B*52:01 | 7 | 14 | 0.70 |
| C*07:02 | B*07:02 | 66 | 37 | 0.69 |
| DRB1*14:54 | DQB1*05:03 | 5 | 11 | 0.67 |
| C*12:02 | DQB1*06:01 | 7 | 8 | 0.66 |
| C*12:02 | DRB1*15:02 | 7 | 8 | 0.66 |
| C*16:01 | B*44:03 | 39 | 48 | 0.65 |
| C*12:03 | B*38:01 | 23 | 10 | 0.65 |
| DRB1*01:01 | DQB1*05:01 | 35 | 85 | 0.62 |
| C*07:01 | B*08:01 | 47 | 19 | 0.61 |

**Supplementary Table 5. Top positively correlated 2-field HLA allele pairs in the East Asian genetic ancestry group (Pearson *r* > 0.6).**

| Allele 1 | Allele 2 | Allele 1 count | Allele 2 count | Pearson *r* |
|---|---|---|---|---|
| DRB1*03:01 | DQB1*02:01 | 33 | 34 | 1.00 |
| DRB1*13:01 | DQB1*06:03 | 5 | 5 | 1.00 |
| DRA*01:03 | DRB1*08:03 | 67 | 68 | 0.98 |
| C*15:05 | B*07:05 | 19 | 18 | 0.98 |
| C*03:02 | B*58:01 | 59 | 56 | 0.95 |
| DRB1*09:01 | DQB1*03:03 | 147 | 162 | 0.95 |
| DPA1*04:01 | DPB1*107:01 | 28 | 24 | 0.91 |
| C*14:03 | DQB1*06:04 | 17 | 20 | 0.88 |
| A*29:01 | B*07:05 | 18 | 18 | 0.87 |
| DRB1*04:05 | DQB1*04:01 | 63 | 48 | 0.87 |
| A*30:01 | B*13:02 | 15 | 15 | 0.86 |
| DRB1*07:01 | DQB1*02:02 | 43 | 30 | 0.85 |
| A*29:01 | C*15:05 | 18 | 19 | 0.85 |
| C*14:02 | B*51:01 | 50 | 58 | 0.80 |
| C*12:02 | B*52:01 | 47 | 36 | 0.78 |
| C*14:03 | B*44:03 | 17 | 26 | 0.77 |
| C*01:02 | B*46:01 | 187 | 144 | 0.76 |
| DPA1*02:07 | DPB1*19:01 | 16 | 15 | 0.76 |
| DRB1*13:02 | DQB1*06:04 | 34 | 20 | 0.75 |
| C*04:03 | B*15:25 | 17 | 13 | 0.73 |
| DRB1*14:05 | DQB1*05:03 | 25 | 51 | 0.70 |
| DRB1*15:01 | DQB1*06:02 | 94 | 51 | 0.69 |
| A*02:07 | B*46:01 | 112 | 144 | 0.68 |
| B*44:03 | DQB1*06:04 | 26 | 20 | 0.67 |
| C*08:01 | B*15:02 | 89 | 48 | 0.67 |
| C*14:03 | DRB1*13:02 | 17 | 34 | 0.66 |
| DRB1*12:02 | DQB1*03:01 | 100 | 182 | 0.66 |
| B*13:02 | DQB1*02:02 | 15 | 30 | 0.65 |
| DRB1*08:03 | DQB1*06:01 | 68 | 132 | 0.65 |
| C*06:02 | B*13:02 | 37 | 15 | 0.65 |
| DRA*01:03 | DQB1*06:01 | 67 | 132 | 0.64 |
| DRB1*16:02 | DQB1*05:02 | 49 | 115 | 0.64 |
| B*07:05 | DRB1*10:01 | 18 | 22 | 0.64 |
| DRB1*13:02 | DQB1*06:09 | 34 | 14 | 0.64 |
| A*33:03 | C*03:02 | 86 | 59 | 0.63 |
| B*58:01 | DQB1*02:01 | 56 | 34 | 0.63 |
| B*07:02 | DRB1*01:01 | 27 | 20 | 0.63 |
| C*15:05 | DRB1*10:01 | 19 | 22 | 0.63 |
| C*08:03 | B*48:01 | 7 | 18 | 0.62 |
| B*58:01 | DRB1*03:01 | 56 | 33 | 0.61 |
| A*01:01 | B*57:01 | 19 | 11 | 0.61 |
| B*13:02 | DPB1*17:01 | 15 | 14 | 0.61 |
| DPA1*02:02 | DPB1*05:01 | 529 | 372 | 0.61 |
| C*03:04 | B*13:01 | 120 | 53 | 0.60 |
| B*13:02 | DRB1*07:01 | 15 | 43 | 0.60 |

**Supplementary Table 6. Top positively correlated 2-field HLA allele pairs in the European genetic ancestry group (Pearson *r* > 0.6).**

| **Allele 1** | **Allele 2** | **Allele 1 count** | **Allele 2 count** | **Pearson *r*** |
|---|---|---|---|---|
| DPA1*02:07 | DPB1*19:01 | 8 | 8 | 1.00 |
| C*17:03 | B*41:02 | 7 | 7 | 1.00 |
| DRB1*03:01 | DQB1*02:01 | 105 | 103 | 0.98 |
| DRB1*15:01 | DQB1*06:02 | 120 | 111 | 0.95 |
| DRB1*11:02 | DQB1*03:19 | 8 | 7 | 0.95 |
| DRB1*13:01 | DQB1*06:03 | 55 | 61 | 0.95 |
| DRB1*08:01 | DQB1*04:02 | 49 | 59 | 0.92 |
| DRB1*13:02 | DQB1*06:04 | 38 | 31 | 0.90 |
| DRB1*14:54 | DQB1*05:03 | 23 | 30 | 0.88 |
| C*07:02 | B*07:02 | 121 | 112 | 0.87 |
| DRB1*01:01 | DQB1*05:01 | 107 | 137 | 0.85 |
| DRB1*07:01 | DQB1*02:02 | 140 | 98 | 0.82 |
| DRB1*16:01 | DQB1*05:02 | 14 | 22 | 0.79 |
| DPA1*01:04 | DPB1*15:01 | 5 | 8 | 0.79 |
| C*03:04 | B*40:01 | 75 | 49 | 0.79 |
| C*16:01 | B*44:03 | 45 | 60 | 0.75 |
| C*05:01 | B*44:02 | 83 | 81 | 0.75 |
| C*08:02 | B*14:02 | 27 | 15 | 0.73 |
| C*07:01 | B*08:01 | 128 | 82 | 0.73 |
| A*29:02 | C*16:01 | 37 | 45 | 0.71 |
| C*08:02 | B*14:01 | 27 | 12 | 0.69 |
| DRB1*04:04 | DQB1*03:02 | 53 | 104 | 0.69 |
| A*66:01 | C*17:03 | 5 | 7 | 0.67 |
| A*66:01 | B*41:02 | 5 | 7 | 0.67 |
| DPA1*02:06 | DPB1*05:01 | 10 | 24 | 0.64 |
| C*04:01 | B*35:01 | 137 | 70 | 0.63 |
| B*08:01 | DRB1*03:01 | 82 | 105 | 0.63 |
| B*08:01 | DQB1*02:01 | 82 | 103 | 0.62 |
| A*01:01 | B*08:01 | 126 | 82 | 0.61 |

| **Supplementary Table 7. Top positively correlated 2-field HLA allele pairs in the South Asian genetic ancestry group (Pearson *r* > 0.6).** | | | | |
|---|---|---|---|---|
| **Allele 1** | **Allele 2** | **Allele 1 count** | **Allele 2 count** | **Pearson *r*** |
| DPA1*04:01 | DPB1*296:01 | 6 | 6 | 1.00 |
| DPA1*01:04 | DPB1*15:01 | 6 | 6 | 1.00 |
| DRB1*03:01 | DQB1*02:01 | 75 | 73 | 0.99 |
| C*03:02 | B*58:01 | 29 | 30 | 0.97 |
| C*08:01 | B*15:02 | 40 | 43 | 0.96 |
| C*15:05 | B*07:05 | 10 | 9 | 0.96 |
| C*07:06 | B*44:03 | 64 | 70 | 0.94 |
| C*04:03 | B*13:01 | 14 | 12 | 0.92 |
| DRB1*13:01 | DQB1*06:03 | 61 | 57 | 0.92 |
| C*07:26 | B*15:25 | 5 | 6 | 0.91 |
| DRB1*14:04 | DQB1*05:03 | 63 | 80 | 0.88 |
| C*12:02 | B*52:01 | 102 | 91 | 0.88 |
| C*07:04 | B*15:18 | 15 | 17 | 0.87 |
| B*35:02 | DRB1*11:04 | 7 | 11 | 0.79 |
| DRA*01:03 | DRB1*08:03 | 21 | 14 | 0.79 |
| C*02:02 | B*27:05 | 6 | 7 | 0.77 |
| C*06:02 | B*57:01 | 140 | 91 | 0.77 |
| C*15:02 | B*40:06 | 100 | 90 | 0.75 |
| DRB1*10:01 | DQB1*05:01 | 51 | 87 | 0.74 |
| DRB1*04:03 | DQB1*03:02 | 41 | 78 | 0.74 |
| DRB1*07:01 | DQB1*02:02 | 190 | 105 | 0.71 |
| A*29:01 | B*07:05 | 9 | 9 | 0.70 |
| DRB1*13:02 | DQB1*06:09 | 23 | 11 | 0.68 |
| C*05:01 | B*44:02 | 9 | 6 | 0.68 |
| C*07:01 | B*15:17 | 26 | 13 | 0.67 |
| A*29:01 | C*15:05 | 9 | 10 | 0.67 |
| A*30:01 | B*13:02 | 8 | 10 | 0.66 |
| DRA*01:02 | DRB1*15:02 | 252 | 120 | 0.64 |
| B*08:01 | DRB1*03:01 | 48 | 75 | 0.64 |
| C*03:04 | B*40:01 | 16 | 14 | 0.64 |
| A*02:03 | B*38:02 | 17 | 16 | 0.64 |
| B*08:01 | DQB1*02:01 | 48 | 73 | 0.64 |
| DRB1*15:06 | DQB1*05:02 | 15 | 36 | 0.63 |
| C*03:03 | B*15:05 | 28 | 12 | 0.62 |
| DRB1*04:02 | DPB1*05:01 | 8 | 8 | 0.62 |
| C*12:04 | B*51:06 | 6 | 19 | 0.62 |
| DRB1*13:02 | DQB1*06:04 | 23 | 9 | 0.62 |

| **Supplementary Table 8. Disjoint graphs (connected components) of HLA alleles across genes**. For this analysis, HLA alleles from different genes were connected by graph edges when the absolute Pearson correlation between the alleles met or exceeded the specified threshold. This procedure was performed separately for each of the five genetic ancestry groups: African (AFR, N = 661), Admixed American (AMR, N = 347), East Asian (EAS, N = 504), European (EUR, N = 503), and South Asian (SAS, N = 489). Connected-component analysis was then applied to each graph to identify maximal connected components, defined as disjoint sets of HLA alleles linked directly or indirectly through correlation-based edges. | | | | | | |
|---|---|---|---|---|---|---|
| **$\|r\|$ threshold** | **Disjoint graphs** | **AFR** | **AMR** | **EAS** | **EUR** | **SAS** |
| 0.3 | N | 26 | 21 | 16 | 21 | 21 |
| | Min, max no. of alleles | [2, 24] | [2, 55] | [2, 57] | [2, 38] | [2, 29] |
| | Mean no. of alleles (SE) | 3.69 (0.86) | 6.00 (2.49) | 6.94 (3.40) | 4.86 (1.70) | 5.38 (1.43) |
| | Min, max no. of genes | [2, 8] | [2, 8] | [2, 8] | [2, 8] | [2, 7] |
| | Mean no. of genes (SE) | 2.58 (0.27) | 3.19 (0.37) | 2.88 (0.44) | 2.67 (0.32) | 2.86 (0.30) |
| 0.4 | N | 32 | 34 | 25 | 27 | 34 |
| | Min, max no. of alleles | [2, 6] | [2, 7] | [2, 19] | [2, 10] | [2, 9] |
| | Mean no. of alleles (SE) | 2.31 (0.15) | 2.91 (0.22) | 3.76 (0.72) | 3.07 (0.35) | 2.97 (0.28) |
| | Min, max no. of genes | [2, 6] | [2, 5] | [2, 6] | [2, 5] | [2, 5] |
| | Mean no. of genes (SE) | 2.16 (0.13) | 2.53 (0.17) | 2.76 (0.27) | 2.52 (0.20) | 2.47 (0.15) |
| 0.5 | N | 28 | 26 | 26 | 26 | 33 |
| | Min, max no. of alleles | [2, 3] | [2, 5] | [2, 9] | [2, 5] | [2, 7] |
| | Mean no. of alleles (SE) | 2.07 (0.05) | 2.46 (0.19) | 2.77 (0.32) | 2.58 (0.20) | 2.45 (0.19) |
| | Min, max no. of genes | [2, 3] | [2, 5] | [2, 6] | [2, 5] | [2, 4] |
| | Mean no. of genes (SE) | 2.04 (0.04) | 2.35 (0.16) | 2.62 (0.23) | 2.42 (0.19) | 2.18 (0.09) |
| 0.6 | N | 21 | 22 | 24 | 21 | 32 |
| | Min, max no. of alleles | [2, 3] | [2, 4] | [2, 6] | [2, 5] | [2, 3] |
| | Mean no. of alleles (SE) | 2.05 (0.05) | 2.09 (0.09) | 2.58 (0.24) | 2.29 (0.16) | 2.09 (0.05) |
| | Min, max no. of genes | [2, 3] | [2, 4] | [2, 6] | [2, 5] | [2, 3] |
| | Mean no. of genes (SE) | 2.05 (0.05) | 2.09 (0.09) | 2.54 (0.23) | 2.24 (0.15) | 2.06 (0.04) |

| **Supplementary Table 9. Principal component analyses on the pairwise Pearson correlation matrix of HLA alleles.** Eigenvalue decomposition was performed on the pairwise Pearson correlation matrix of HLA alleles. Only HLA alleles with minor allele count (MAC) ≥5 were used. $M_{eff}^{eigen}$ - eigenvalue-based effective number of independent tests. Genetic ancestry groups: All combined (ALL, N=2,504) African (AFR, N = 661), Admixed American (AMR, N = 347), East Asian (EAS, N = 504), European (EUR, N = 503), and South Asian (SAS, N = 489). | | | | |
|---|---|---|---|---|
| **Group** | **N alleles (MAC ≥5)** | **PCs (λ > 1)** | $\boldsymbol{M_{eff}^{eigen}}$ | **Δ $\boldsymbol{M_{eff}^{eigen}}$ vs ALL** |
| *Main analyses* | | | | |
| ALL | 283 | 116 | 202 | -- |
| AFR | 148 | 58 | 98 | -- |
| AMR | 153 | 57 | 91 | -- |
| EAS | 135 | 48 | 79 | -- |
| EUR | 133 | 51 | 82 | -- |
| SAS | 138 | 53 | 82 | -- |
| *Leave-one-out analyses* | | | | |
| No AFR | 245 | 99 | 168 | −34 |
| No AMR | 261 | 106 | 184 | −19 |
| No EAS | 256 | 105 | 179 | −23 |
| No EUR | 270 | 110 | 190 | −12 |
| No SAS | 264 | 106 | 185 | −17 |

| **Supplementary Table 10. Eigenvalue-based effective number of independent tests by minor allele count.** Eigenvalue decomposition was performed on the pairwise Pearson correlation matrix of HLA alleles. Genetic ancestry groups: All combined (ALL, N=2,504) African (AFR, N = 661), Admixed American (AMR, N = 347), East Asian (EAS, N = 504), European (EUR, N = 503), and South Asian (SAS, N = 489). | | | | | | |
|---|---|---|---|---|---|---|
| | **MAC≥5** | | **MAC≥10** | | **MAC≥20** | |
| **Dataset** | **N alleles** | $\boldsymbol{M_{eff}^{eigen}}$ **(% total)** | **N alleles** | $\boldsymbol{M_{eff}^{eigen}}$ **(% total)** | **N_alleles** | $\boldsymbol{M_{eff}^{eigen}}$**(% total)** |
| ALL | 283 | 202 (71) | 236 | 165 (70) | 196 | 135 (69) |
| AFR | 148 | 98 (66) | 128 | 84 (65) | 103 | 68 (66) |
| AMR | 153 | 91 (59) | 109 | 67 (61) | 64 | 40 (63) |
| EAS | 135 | 79 (59) | 110 | 64 (58) | 84 | 50 (60) |
| EUR | 133 | 82 (61) | 105 | 64 (61) | 77 | 47 (61) |
| SAS | 138 | 82 (60) | 107 | 66 (61) | 77 | 48 (62) |

| **Supplementary Table 11. Simulation-based effective number of independent tests by minor allele count using linear regression model.** The results are based on 30,000 simulated continuous phenotypes with fixed effects from sex and top 10 principal components from genetic relatedness matrix. Genetic ancestry groups: All combined (ALL, N=2,504) African (AFR, N = 661), Admixed American (AMR, N = 347), East Asian (EAS, N = 504), European (EUR, N = 503), and South Asian (SAS, N = 489). | | | | | | | | |
|---|---|---|---|---|---|---|---|---|
| | **MAC≥5** | | **MAC≥10** | | **MAC≥20** | | **MAC≥50** | |
| **Dataset** | **N alleles** | $M_{eff}^{perm}$ **(% total)** | **N alleles** | $M_{eff}^{perm}$ **(% total)** | **N alleles** | $M_{eff}^{perm}$ **(% total)** | **N alleles** | $M_{eff}^{perm}$ **(% total)** |
| ALL | 283 | 260 (92) | 236 | 216 (91) | 196 | 182 (93) | 143 | 130 (91) |
| AFR | 148 | 134 (91) | 128 | 116 (90) | 103 | 93 (90) | 62 | 54 (88) |
| AMR | 153 | 142 (93) | 109 | 104 (96) | 64 | 58 (90) | 21 | 20 (95) |
| EAS | 135 | 124 (92) | 110 | 100 (91) | 84 | 75 (89) | 44 | 39 (89) |
| EUR | 133 | 117 (88) | 105 | 93 (89) | 77 | 66 (86) | 43 | 36 (85) |
| SAS | 138 | 128 (92) | 107 | 103 (96) | 77 | 70 (91) | 46 | 43 (93) |

| **Supplementary Table 12. Simulation-based effective number of independent tests by minor allele count using Firth logistic regression model.** | | | | | | | | | |
|---|---|---|---|---|---|---|---|---|---|
| | | **MAC≥5** | | **MAC≥10** | | **MAC≥20** | | **MAC≥50** | |
| **Dataset** | **% Cases** | **N alleles** | $M_{eff}^{sim}$ **(% total)** | **N alleles** | $M_{eff}^{sim}$ **(% total)** | **N alleles** | $M_{eff}^{sim}$ **(% total)** | **N alleles** | $M_{eff}^{sim}$ **(% total)** |
| ALL | 10 | 283 | 223 (79) | 236 | 187 (79) | 196 | 159 (81) | 143 | 124 (87) |
| AFR | 10 | 148 | 108 (73) | 128 | 95 (74) | 103 | 78 (75) | 62 | 53 (85) |
| AMR | 10 | 153 | 94 (62) | 109 | 70 (64) | 64 | 42 (65) | 21 | 18 (84) |
| EAS | 10 | 135 | 93 (69) | 110 | 74 (67) | 84 | 59 (70) | 44 | 33 (76) |
| EUR | 10 | 133 | 90 (68) | 105 | 71 (68) | 77 | 54 (70) | 43 | 33 (76) |
| SAS | 10 | 138 | 87 (63) | 107 | 72 (67) | 77 | 52 (67) | 46 | 33 (72) |
| ALL | 20 | 283 | 211 (75) | 236 | 188 (80) | 196 | 167 (85) | 143 | 132 (92) |
| AFR | 20 | 148 | 108 (73) | 128 | 96 (75) | 103 | 81 (79) | 62 | 54 (87) |
| AMR | 20 | 153 | 97 (63) | 109 | 75 (69) | 64 | 48 (75) | 21 | 18 (87) |
| EAS | 20 | 135 | 92 (68) | 110 | 77 (70) | 84 | 63 (75) | 44 | 36 (82) |
| EUR | 20 | 133 | 89 (67) | 105 | 73 (70) | 77 | 58 (75) | 43 | 35 (82) |
| SAS | 20 | 138 | 94 (68) | 107 | 81 (75) | 77 | 61 (79) | 46 | 40 (86) |

| **Supplementary Table 13. Number of distinct 2-field and 4-field alleles in classical HLA genes typed in the 1000 Genomes Project's long read sequencing data.** Results are based on 415 individuals representing different genetic ancestries after quality control. MAC – minor allele count. | | | | | | |
|---|---|---|---|---|---|---|
| | **MAC≥1** | | | **MAC≥5** | | |
| **Gene (class)** | **2-field** | **4-field** | **Fold increase** | **2-field** | **4-field** | **Fold increase** |
| *A (Class I)* | 41 | 59 | 1.4 | 26 | 28 | 1.1 |
| *B (Class I)* | 58 | 96 | 1.7 | 43 | 53 | 1.2 |
| *C (Class I)* | 28 | 65 | 2.3 | 24 | 36 | 1.5 |
| *DPA1 (Class II)* | 7 | 27 | 3.9 | 5 | 14 | 2.8 |
| *DPB1 (Class II)* | 31 | 100 | 3.2 | 18 | 25 | 1.4 |
| *DQA1 (Class II)* | 14 | 33 | 2.4 | 13 | 26 | 2.0 |
| *DQB1 (Class II)* | 17 | 59 | 3.5 | 16 | 33 | 2.1 |
| *DRA (Class II)* | 2 | 28 | 14.0 | 2 | 22 | 11.0 |
| *DRB1 (Class II)* | 38 | 109 | 2.9 | 28 | 36 | 1.3 |
| **Total** | **236** | **576** | **2.4** | **175** | **273** | **1.6** |

| **Supplementary Table 14. Top positively correlated 2-field HLA allele pairs typed in the 1000 Genomes Project's long read sequencing data. (Pearson *r* > 0.4).** Results are based on 415 individuals representing different genetic ancestries after quality control. | | | | |
|---|---|---|---|---|
| **Allele 1** | **Allele 2** | **Allele 1 count** | **Allele 2 count** | **Pearson *r*** |
| C*12:02 | B*52:01 | 19 | 17 | 0.95 |
| C*08:02 | B*14:02 | 16 | 14 | 0.95 |
| DRB1*07:01 | DQA1*02:01 | 100 | 104 | 0.93 |
| DQA1*04:01 | DQB1*04:02 | 54 | 59 | 0.89 |
| DRB1*13:01 | DQB1*06:03 | 29 | 26 | 0.86 |
| DRB1*10:01 | DQA1*01:05 | 20 | 18 | 0.85 |
| DQA1*05:01 | DQB1*02:01 | 55 | 43 | 0.83 |
| DQA1*03:01 | DQB1*03:02 | 90 | 83 | 0.83 |
| DRB1*01:01 | DQA1*01:01 | 39 | 62 | 0.82 |
| C*17:01 | B*42:01 | 23 | 14 | 0.81 |
| C*15:05 | B*07:05 | 8 | 5 | 0.79 |
| DQA1*02:01 | DQB1*02:02 | 104 | 80 | 0.78 |
| DQA1*01:01 | DQB1*05:01 | 62 | 97 | 0.75 |
| A*29:01 | B*07:05 | 6 | 5 | 0.73 |
| C*03:02 | B*58:01 | 22 | 33 | 0.72 |
| DRB1*03:01 | DQB1*02:01 | 57 | 43 | 0.71 |
| A*02:07 | B*46:01 | 11 | 19 | 0.71 |
| DRB1*11:02 | DQB1*03:19 | 6 | 10 | 0.71 |
| DRB1*04:03 | DQB1*03:02 | 50 | 83 | 0.71 |
| DRB1*07:01 | DQB1*02:02 | 100 | 80 | 0.71 |
| DRB1*04:03 | DQA1*03:01 | 50 | 90 | 0.69 |
| DPA1*02:02 | DPB1*05:01 | 128 | 72 | 0.69 |
| DRB1*09:01 | DQA1*03:02 | 81 | 40 | 0.68 |
| DRB1*12:02 | DQA1*06:01 | 11 | 20 | 0.67 |
| C*05:01 | B*44:02 | 37 | 28 | 0.67 |
| DQA1*03:02 | DQB1*03:03 | 40 | 74 | 0.67 |
| DQA1*01:04 | DQB1*05:03 | 46 | 36 | 0.67 |
| C*02:02 | B*27:05 | 19 | 16 | 0.66 |
| DQA1*05:05 | DQB1*03:01 | 75 | 112 | 0.65 |
| DRB1*04:05 | DQB1*04:01 | 12 | 7 | 0.65 |
| DRB1*03:01 | DQA1*05:01 | 57 | 55 | 0.63 |
| B*08:01 | DQB1*02:01 | 34 | 43 | 0.63 |
| DRB1*14:01 | DQA1*01:04 | 22 | 46 | 0.63 |
| C*06:02 | B*57:01 | 72 | 33 | 0.62 |
| A*30:01 | C*17:01 | 24 | 23 | 0.62 |
| DRB1*01:01 | DQB1*05:01 | 39 | 97 | 0.62 |
| C*03:03 | B*15:01 | 29 | 26 | 0.62 |
| DRB1*08:02 | DQA1*04:01 | 36 | 54 | 0.61 |
| DRB1*13:02 | DQB1*06:09 | 38 | 21 | 0.61 |
| DQA1*01:03 | DQB1*06:01 | 80 | 60 | 0.6 |
| C*04:01 | B*35:01 | 116 | 70 | 0.6 |
| DQA1*01:02 | DQB1*06:02 | 157 | 68 | 0.59 |
| C*12:03 | B*38:01 | 25 | 9 | 0.59 |
| A*30:01 | B*42:01 | 24 | 14 | 0.59 |
| DRB1*08:02 | DQB1*04:02 | 36 | 59 | 0.58 |
| A*29:01 | C*15:05 | 6 | 8 | 0.57 |
| C*08:01 | B*15:02 | 26 | 14 | 0.57 |
| C*07:18 | B*58:01 | 17 | 33 | 0.57 |
| B*08:01 | DQA1*05:01 | 34 | 55 | 0.55 |
| A*33:03 | C*03:02 | 33 | 22 | 0.54 |
| DRB1*15:01 | DQB1*06:02 | 88 | 68 | 0.54 |
| C*15:02 | B*40:06 | 38 | 20 | 0.53 |
| DRB1*15:01 | DQA1*01:02 | 88 | 157 | 0.53 |
| B*08:01 | DRB1*03:01 | 34 | 57 | 0.53 |
| C*01:02 | B*46:01 | 58 | 19 | 0.53 |
| DQA1*01:03 | DQB1*06:03 | 80 | 26 | 0.5 |
| B*15:21 | DQB1*06:04 | 11 | 17 | 0.49 |
| DRB1*11:02 | DPB1*131:01 | 6 | 8 | 0.49 |
| C*07:02 | B*07:02 | 85 | 53 | 0.49 |
| C*08:01 | B*48:01 | 26 | 13 | 0.49 |
| DRB1*11:01 | DQA1*05:05 | 52 | 75 | 0.49 |
| C*03:04 | B*40:01 | 58 | 36 | 0.48 |

| | | | | |
|---|---|---|---|---|
| DRB1*13:02 | DQB1*06:04 | 38 | 17 | 0.47 |
| DRB1*01:02 | DQA1*01:01 | 14 | 62 | 0.47 |
| A*02:11 | B*40:06 | 17 | 20 | 0.47 |
| DRB1*13:01 | DQA1*01:03 | 29 | 80 | 0.47 |
| B*58:01 | DQB1*06:09 | 33 | 21 | 0.46 |
| C*07:06 | B*44:03 | 14 | 57 | 0.46 |
| DRB1*16:01 | DQB1*05:02 | 12 | 36 | 0.46 |
| A*29:02 | C*16:01 | 22 | 41 | 0.46 |
| C*07:01 | B*08:01 | 54 | 34 | 0.46 |
| B*15:02 | DQA1*06:01 | 14 | 20 | 0.46 |
| A*33:03 | B*58:01 | 33 | 33 | 0.45 |
| C*07:01 | B*15:21 | 54 | 11 | 0.45 |
| C*16:01 | B*44:03 | 41 | 57 | 0.44 |
| DRB1*09:01 | DQB1*03:03 | 81 | 74 | 0.44 |
| A*01:01 | B*08:01 | 67 | 34 | 0.44 |
| DPA1*01:03 | DPB1*04:01 | 485 | 181 | 0.43 |
| A*02:07 | C*01:02 | 11 | 58 | 0.43 |
| B*15:02 | DRB1*12:02 | 14 | 11 | 0.43 |
| DRB1*03:02 | DQA1*04:01 | 6 | 54 | 0.42 |
| C*14:02 | B*51:06 | 19 | 9 | 0.41 |
| A*01:01 | B*57:01 | 67 | 33 | 0.41 |
| B*42:01 | DRB1*03:02 | 14 | 6 | 0.41 |
| C*14:02 | B*51:01 | 19 | 49 | 0.41 |

**Supplementary Table 15. Top positively correlated 4-field HLA allele pairs typed in the 1000 Genomes Project's long read sequencing data (Pearson *r* > 0.5).** Results are based on 415 individuals representing different genetic ancestries after quality control.

| Allele 1 | Allele 2 | Allele 1 count | Allele 2 count | Pearson *r* |
|---|---|---|---|---|
| DRA*01:02:02:14 | DQA1*06:01:01:02 | 13 | 20 | 0.96 |
| C*03:02:02:05 | B*58:01:01:03 | 22 | 19 | 0.95 |
| DRA*01:01:01:11 | DQA1*01:05:01:01 | 16 | 18 | 0.94 |
| C*12:02:02:01 | B*52:01:01:02 | 17 | 17 | 0.91 |
| C*08:02:01:01 | B*14:02:01:01 | 15 | 14 | 0.91 |
| DQA1*01:03:01:02 | DQB1*06:03:01:01 | 25 | 23 | 0.89 |
| DRA*01:02:02:04 | DQB1*06:09:01:01 | 18 | 21 | 0.89 |
| C*07:18:01:01 | B*58:01:01:01 | 17 | 14 | 0.88 |
| DRB1*13:01:01:01 | DQA1*01:03:01:02 | 26 | 25 | 0.88 |
| C*06:02:01:02 | B*50:01:01:01 | 9 | 12 | 0.86 |
| DRA*01:01:01:13 | DQA1*01:01:02:01 | 13 | 9 | 0.84 |
| DPA1*01:03:01:05 | DPB1*04:02:01:02 | 85 | 97 | 0.83 |
| DQA1*02:01:01:01 | DQB1*02:02:01:01 | 104 | 74 | 0.83 |
| DQA1*01:01:02:01 | DQB1*05:01:01:01 | 9 | 13 | 0.83 |
| DQA1*05:01:01:01 | DQB1*02:01:01:01 | 32 | 43 | 0.82 |
| DQA1*01:01:01:01 | DQB1*05:01:01:03 | 44 | 53 | 0.81 |
| DRB1*13:01:01:01 | DQB1*06:03:01:01 | 26 | 23 | 0.81 |
| DRB1*01:01:01:01 | DQB1*05:01:01:03 | 38 | 53 | 0.81 |
| C*17:01:01:02 | B*42:01:01:01 | 23 | 13 | 0.8 |
| DQA1*03:02:01:01 | DQB1*03:03:02:02 | 32 | 40 | 0.8 |
| DRB1*07:01:01:01 | DQA1*02:01:01:01 | 75 | 104 | 0.8 |
| DRB1*01:01:01:01 | DQA1*01:01:01:01 | 38 | 44 | 0.8 |
| C*15:05:02:01 | B*07:05:01:01 | 8 | 5 | 0.79 |
| DRA*01:01:01:11 | DRB1*10:01:01:03 | 16 | 16 | 0.77 |
| DQA1*04:01:01:02 | DQB1*04:02:01:04 | 44 | 41 | 0.76 |
| DRA*01:01:01:08 | DQB1*03:19:01:01 | 10 | 10 | 0.76 |
| C*14:02:01:01 | B*51:01:01:01 | 19 | 13 | 0.75 |
| DPA1*02:07:01:04 | DPB1*04:01:01:41 | 9 | 7 | 0.75 |
| C*16:01:01:01 | B*44:03:01:01 | 41 | 21 | 0.75 |
| C*07:04:01:01 | DQB1*06:02:01:04 | 6 | 6 | 0.75 |
| DRA*01:01:01:11 | DQB1*05:01:01:05 | 16 | 10 | 0.74 |
| DPA1*01:03:01:03 | DPB1*03:01:01:01 | 45 | 33 | 0.74 |
| DPA1*01:03:01:01 | DPB1*02:01:02:01 | 105 | 121 | 0.73 |
| DRB1*01:02:01:01 | DQA1*01:01:02:01 | 13 | 9 | 0.73 |
| DRB1*07:01:01:01 | DQB1*02:02:01:01 | 75 | 74 | 0.73 |
| C*07:01:01:01 | B*08:01:01:01 | 28 | 25 | 0.73 |
| DRB1*10:01:01:03 | DQA1*01:05:01:01 | 16 | 18 | 0.73 |
| A*29:01:01:01 | B*07:05:01:01 | 6 | 5 | 0.73 |
| DPA1*01:03:01:34 | DPB1*104:01:01:02 | 15 | 8 | 0.72 |
| C*07:02:01:03 | B*07:02:01:01 | 27 | 41 | 0.72 |
| C*07:01:02:01 | B*15:21:01:01 | 14 | 11 | 0.71 |
| A*02:07:01:01 | B*46:01:01:01 | 11 | 17 | 0.71 |
| DRA*01:02:02:04 | DRB1*13:02:01:01 | 18 | 30 | 0.7 |
| DRB1*12:02:01:01 | DQB1*03:01:01:22 | 8 | 5 | 0.7 |
| DQA1*01:05:01:01 | DQB1*05:01:01:05 | 18 | 10 | 0.7 |
| DRA*01:02:02:14 | DRB1*12:02:01:01 | 13 | 8 | 0.69 |
| DRA*01:01:01:13 | DQB1*05:01:01:01 | 13 | 13 | 0.69 |
| C*06:02:01:01 | B*57:01:01:01 | 60 | 32 | 0.68 |
| C*05:01:01:02 | B*44:02:01:01 | 21 | 22 | 0.68 |
| B*35:02:01:02 | DQB1*03:01:01:02 | 7 | 5 | 0.67 |
| DRB1*13:02:01:01 | DQA1*01:02:01:04 | 30 | 41 | 0.67 |
| DQA1*03:01:01:01 | DQB1*03:02:01:01 | 90 | 60 | 0.67 |
| C*08:01:01:01 | B*15:02:01:01 | 24 | 10 | 0.66 |
| B*37:01:01:01 | DQB1*05:01:01:05 | 8 | 10 | 0.66 |
| DQA1*01:03:01:01 | DQB1*06:01:01:01 | 44 | 60 | 0.66 |
| DRA*01:02:02:14 | DQB1*03:01:01:22 | 13 | 5 | 0.64 |
| DRB1*13:02:01:01 | DQB1*06:09:01:01 | 30 | 21 | 0.63 |
| C*03:03:01:01 | B*15:01:01:01 | 26 | 25 | 0.63 |
| DRA*01:01:01:13 | DRB1*01:02:01:01 | 13 | 13 | 0.63 |
| DQA1*05:05:01:01 | DQB1*03:01:01:03 | 58 | 34 | 0.62 |
| C*02:02:02:01 | B*27:05:02:01 | 18 | 16 | 0.62 |
| DRA*01:02:02:11 | DQB1*04:02:01:04 | 26 | 41 | 0.62 |

| | | | | |
|---|---|---|---|---|
| DQA1*01:02:01:04 | DQB1*06:04:01:01 | 41 | 17 | 0.61 |
| DQA1*01:02:01:04 | DQB1*06:09:01:01 | 41 | 21 | 0.61 |
| DRB1*03:01:01:01 | DQB1*02:01:01:01 | 37 | 43 | 0.61 |
| B*18:01:01:81 | DQB1*06:02:01:04 | 10 | 6 | 0.61 |
| C*07:04:01:01 | B*18:01:01:81 | 6 | 10 | 0.61 |
| DRB1*12:01:01:03 | DQA1*05:05:01:03 | 8 | 13 | 0.61 |
| DRB1*04:03:01:01 | DQA1*03:01:01:01 | 39 | 90 | 0.61 |
| DRB1*01:02:01:01 | DQB1*05:01:01:01 | 13 | 13 | 0.6 |
| DRA*01:02:02:11 | DRB1*08:02:01:01 | 26 | 36 | 0.6 |
| DRB1*15:01:01:02 | DQB1*06:02:01:01 | 65 | 57 | 0.6 |
| C*07:06:01:01 | B*44:03:01:10 | 14 | 33 | 0.6 |
| C*16:02:01:01 | B*51:01:01:09 | 5 | 11 | 0.6 |
| DRA*01:02:02:04 | DQA1*01:02:01:04 | 18 | 41 | 0.59 |
| DRA*01:01:01:05 | DQB1*05:01:01:03 | 40 | 53 | 0.59 |
| DQA1*05:05:01:03 | DQB1*03:01:01:05 | 13 | 8 | 0.59 |
| C*12:03:01:01 | B*38:01:01:01 | 25 | 9 | 0.59 |
| A*33:03:01:01 | B*58:01:01:03 | 33 | 19 | 0.59 |
| DPA1*01:03:01:04 | DPB1*04:01:01:01 | 69 | 163 | 0.59 |
| C*01:02:01:01 | B*46:01:01:01 | 48 | 17 | 0.58 |
| DRA*01:02:02:10 | DQA1*01:03:01:01 | 20 | 44 | 0.58 |
| DRA*01:02:02:09 | DQB1*03:01:01:05 | 13 | 8 | 0.58 |
| DPA1*02:01:08:03 | DPB1*01:01:01:01 | 19 | 32 | 0.57 |
| C*15:02:01:01 | B*40:06:01:12 | 38 | 20 | 0.57 |
| DRA*01:02:02:14 | DQB1*03:01:01:12 | 13 | 5 | 0.57 |
| A*29:01:01:01 | C*15:05:02:01 | 6 | 8 | 0.57 |
| DRA*01:01:01:04 | DQB1*05:01:01:03 | 20 | 53 | 0.56 |
| B*15:02:01:01 | DQB1*03:01:01:22 | 10 | 5 | 0.56 |
| DRB1*12:02:01:01 | DQA1*06:01:01:02 | 8 | 20 | 0.56 |
| B*37:01:01:01 | DRA*01:01:01:11 | 8 | 16 | 0.55 |
| A*30:01:01:01 | C*17:01:01:02 | 22 | 23 | 0.55 |
| DPA1*02:02:02:01 | DPB1*05:01:01:01 | 124 | 47 | 0.55 |
| DRB1*12:01:01:03 | DQB1*03:01:01:05 | 8 | 8 | 0.55 |
| A*33:03:01:01 | C*03:02:02:05 | 33 | 22 | 0.54 |
| DQA1*03:03:01:03 | DQB1*04:01:01:03 | 6 | 5 | 0.54 |
| DRA*01:02:02:06 | DRB1*13:03:01:01 | 6 | 5 | 0.54 |
| DRB1*08:02:01:01 | DQA1*04:01:01:02 | 36 | 44 | 0.54 |
| A*29:02:01:01 | B*44:03:01:01 | 15 | 21 | 0.54 |
| DRB1*10:01:01:03 | DQB1*05:01:01:02 | 16 | 13 | 0.53 |
| DRB1*09:01:02:01 | DQB1*03:03:02:02 | 39 | 40 | 0.53 |
| B*57:01:01:01 | DQB1*03:03:02:01 | 32 | 32 | 0.53 |
| DPA1*01:03:01:02 | DPB1*04:01:01:01 | 129 | 163 | 0.53 |
| B*08:01:01:02 | DQA1*05:01:01:01 | 9 | 32 | 0.53 |
| DRA*01:01:01:05 | DRB1*01:01:01:01 | 40 | 38 | 0.52 |
| DRB1*03:01:01:03 | DQA1*05:01:01:01 | 6 | 32 | 0.52 |
| DRA*01:02:02:10 | DRB1*15:02:01:02 | 20 | 12 | 0.52 |
| DRA*01:01:01:05 | DQA1*01:01:01:01 | 40 | 44 | 0.52 |
| DRA*01:01:01:11 | DQB1*05:01:01:02 | 16 | 13 | 0.52 |
| A*30:01:01:01 | B*42:01:01:01 | 22 | 13 | 0.52 |
| C*07:01:02:01 | DQB1*06:04:01:01 | 14 | 17 | 0.51 |
| A*02:07:01:01 | DQA1*03:02:01:02 | 11 | 8 | 0.51 |
| DRA*01:01:01:04 | DRB1*01:01:01:01 | 20 | 38 | 0.51 |
| C*03:04:01:02 | B*13:01:01:11 | 32 | 5 | 0.51 |
| DQA1*06:01:01:02 | DQB1*03:01:01:22 | 20 | 5 | 0.5 |
| DQA1*06:01:01:02 | DQB1*03:01:01:12 | 20 | 5 | 0.5 |
| DRA*01:02:02:01 | DQA1*05:01:01:02 | 48 | 23 | 0.5 |

**Supplementary Table 16. Eigenvalue-based effective number of independent tests by minor allele count for 2-field and 4-field HLA alleles typed in the 1000 Genomes Project's long read sequencing data.** Results are based on 415 individuals representing different genetic ancestries after quality control.

| | **2-field** | | | **4-field** | | |
|---|---|---|---|---|---|---|
| **MAC** | **N alleles** | **PCs (λ > 1)** | $M_{eff}^{eigen}$ **(% total)** | **N alleles** | **PCs (λ > 1)** | $M_{eff}^{eigen}$ **(% total)** |
| ≥5 | 175 | 64 | 105 (60) | 273 | 96 | 150 (55) |
| ≥10 | 137 | 50 | 82 (60) | 183 | 63 | 105 (57) |
| ≥20 | 94 | 34 | 57 (60) | 109 | 37 | 66 (60) |
| ≥50 | 46 | 17 | 29 (62) | 28 | 11 | 19 (68) |

**Supplementary Table 17. Simulation-based effective number of independent tests by minor allele count using liner regression model for 2-field and 4-field HLA alleles typed in the 1000 Genomes Project's long read sequencing data.** Results are based on 415 individuals representing different genetic ancestries after quality control.

| | 2-field | | 4-field | |
|---|---|---|---|---|
| **MAC** | **N alleles** | $M_{eff}^{sim}$ **(% total)** | **N alleles** | $M_{eff}^{sim}$ **(% total)** |
| ≥5 | 175 | 164 (94) | 273 | 243 (89) |
| ≥10 | 137 | 124 (91) | 183 | 167 (91) |
| ≥20 | 94 | 86 (91) | 109 | 101 (92) |
| ≥50 | 46 | 41 (89) | 28 | 25 (89) |

### 4. Supplementary Figures

**A** **B**

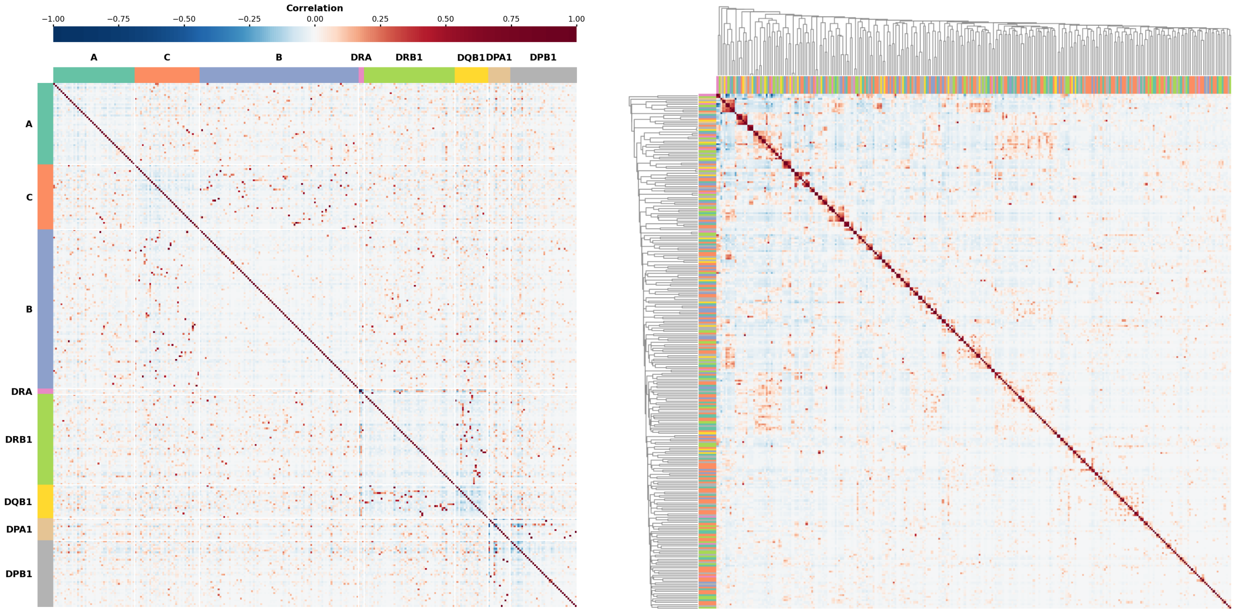


**Supplementary Figure 1. Correlation between classical 2-field HLA alleles.** Each heatmap cell indicates the Pearson correlation coefficient between two alleles. Blue cells indicate negative Pearson correlation coefficients, red cells indicate positive Pearson correlation coefficients, and white cells indicate zero or near-zero correlation between allele pairs. **A.** Alleles are grouped by HLA gene, indicated by the different colors of the track. **B**. The same heatmap but after hierarchical clustering; alleles belonging to the same gene have the same color as in the left panel. The complete reshuffle of the colors in the track means that strong correlations are observed between alleles from different HLA genes.

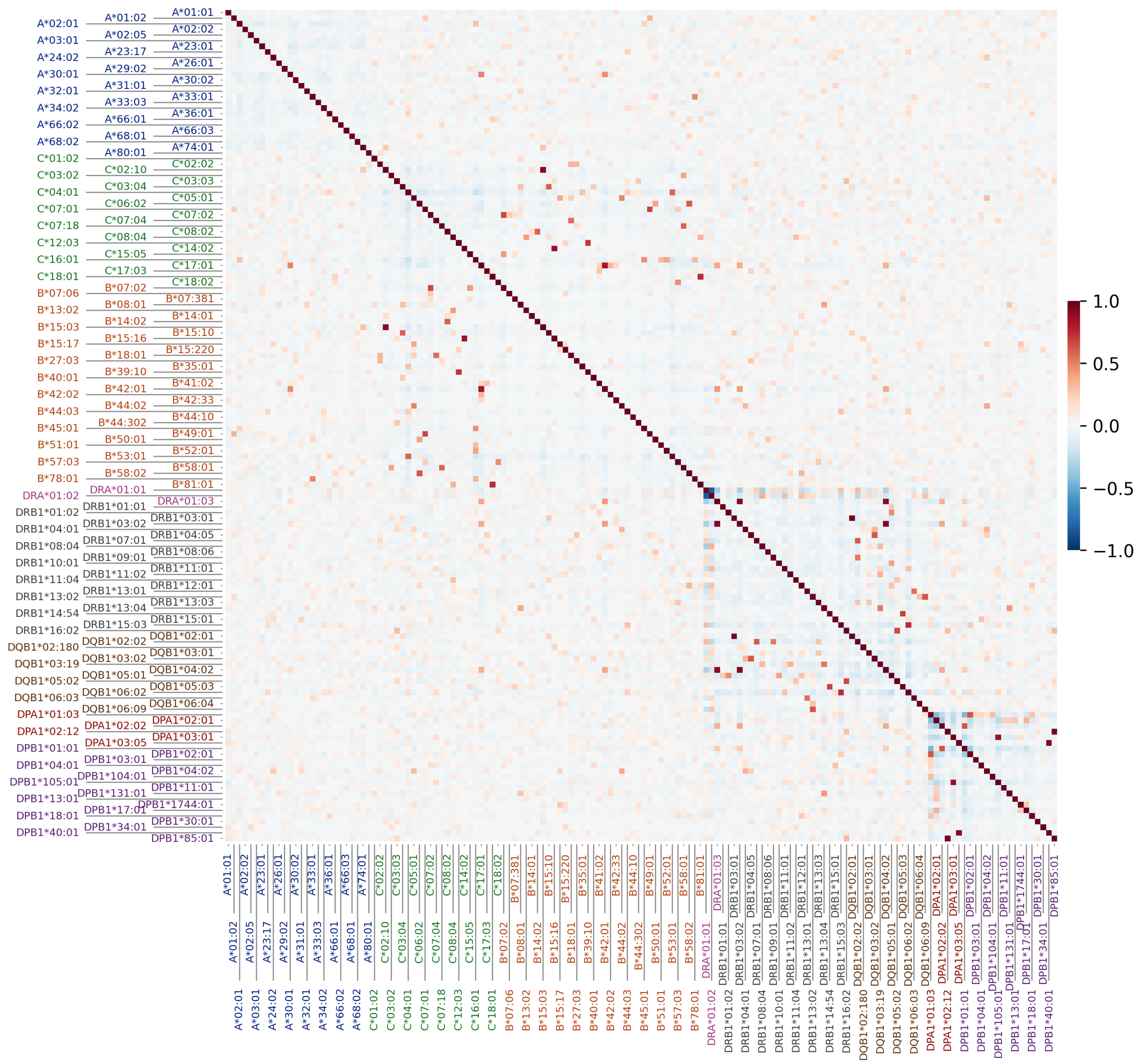


**Supplementary Figure 2A. Correlation between classical 2-field HLA alleles in the African genetic ancestry group.** Blue cells indicate negative Pearson correlation coefficients, red cells indicate positive Pearson correlation coefficients, and white cells indicate zero or near-zero correlation between allele pairs. Alleles are ordered alphanumerically by name. Alleles belonging to the same HLA gene are shown in the same color.

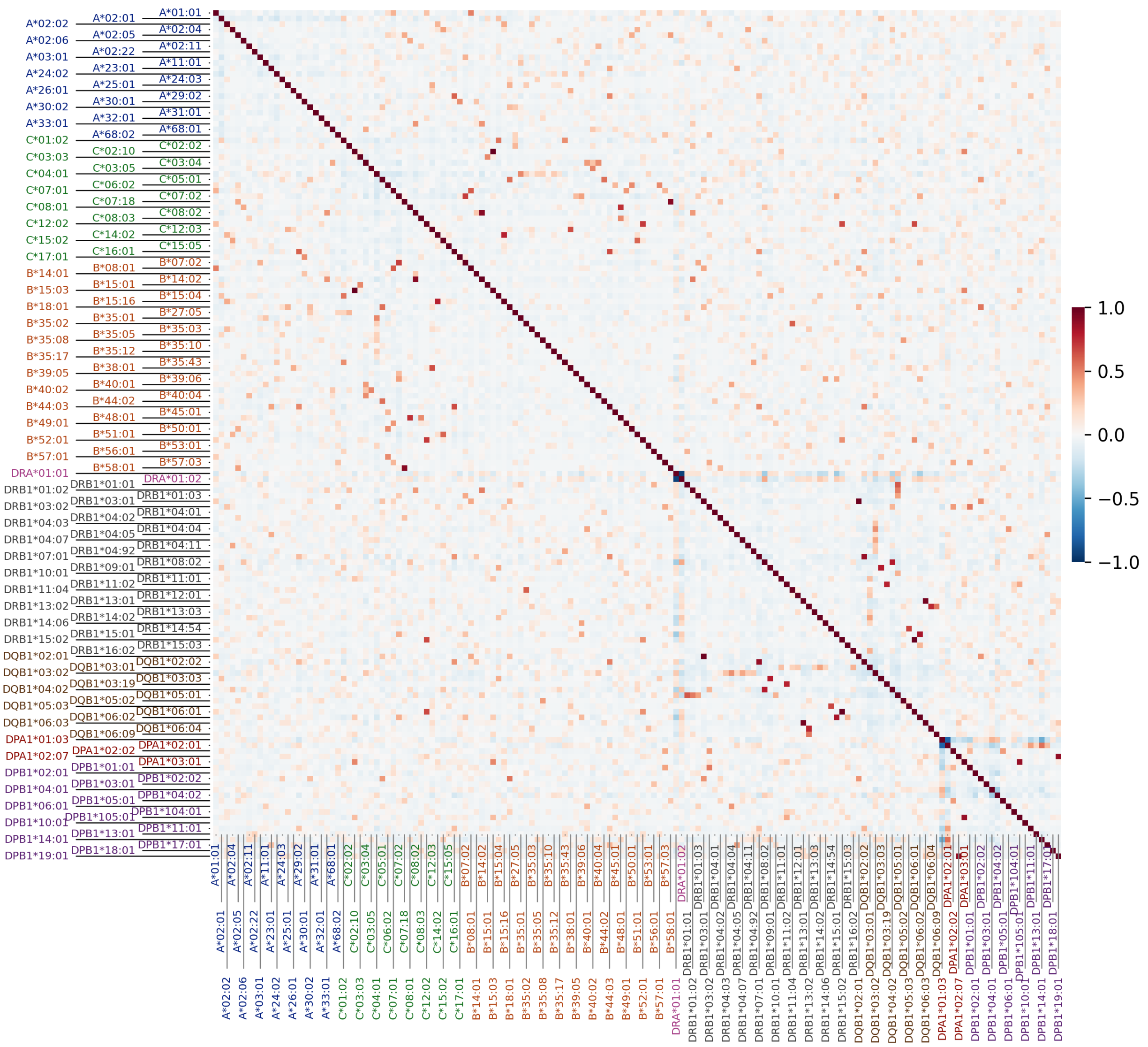


**Supplementary Figure 2B. Correlation between classical 2-field HLA alleles in the Admixed American genetic ancestry group.** Blue cells indicate negative Pearson correlation coefficients, red cells indicate positive Pearson correlation coefficients, and white cells indicate zero or near-zero correlation between allele pairs. Alleles are ordered alphanumerically by name. Alleles belonging to the same HLA gene are shown in the same color.

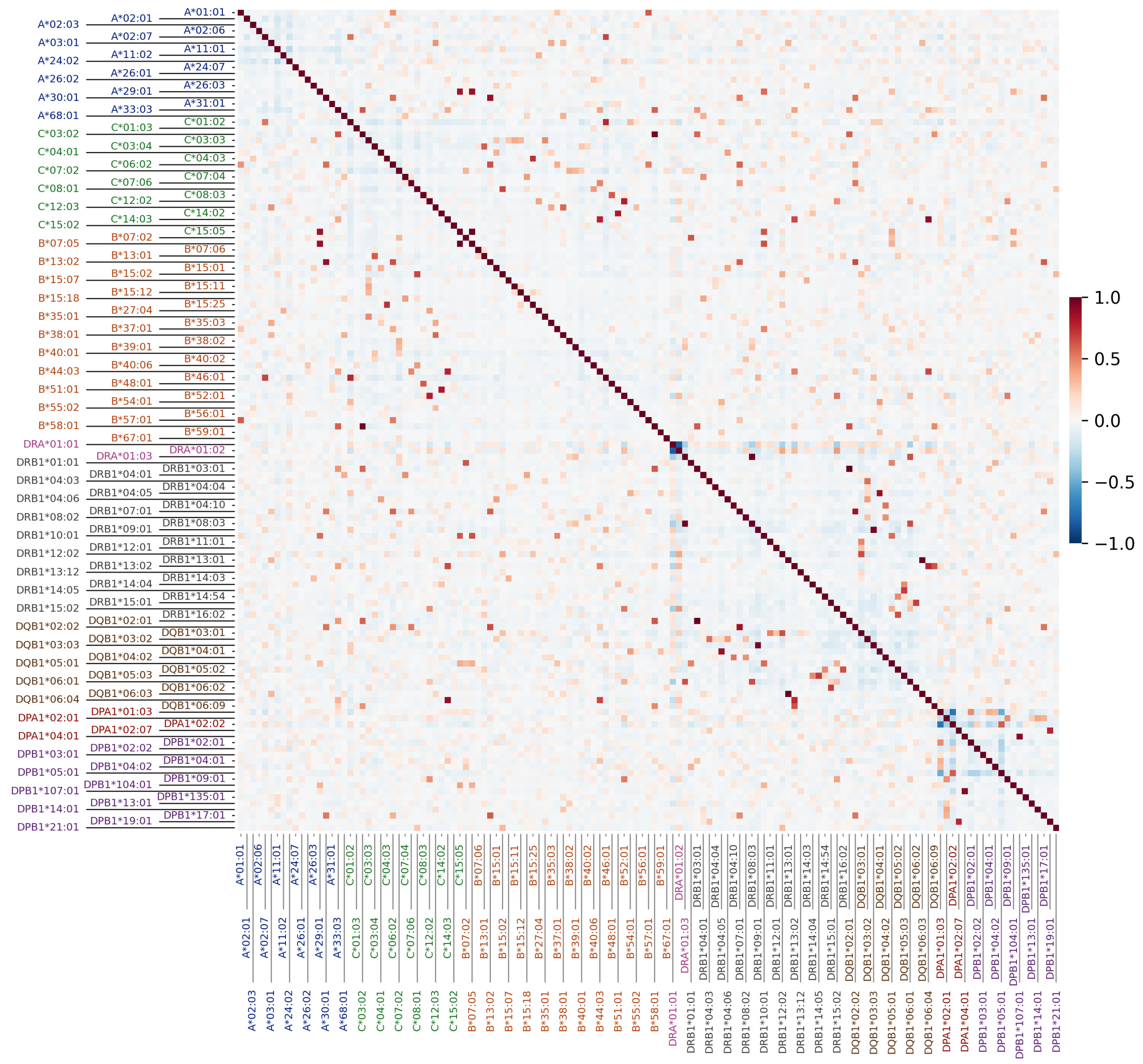


**Supplementary Figure 2C. Correlation between classical 2-field HLA alleles in the East Asian genetic ancestry group.** Blue cells indicate negative Pearson correlation coefficients, red cells indicate positive Pearson correlation coefficients, and white cells indicate zero or near-zero correlation between allele pairs. Alleles are ordered alphanumerically by name. Alleles belonging to the same HLA gene are shown in the same color.

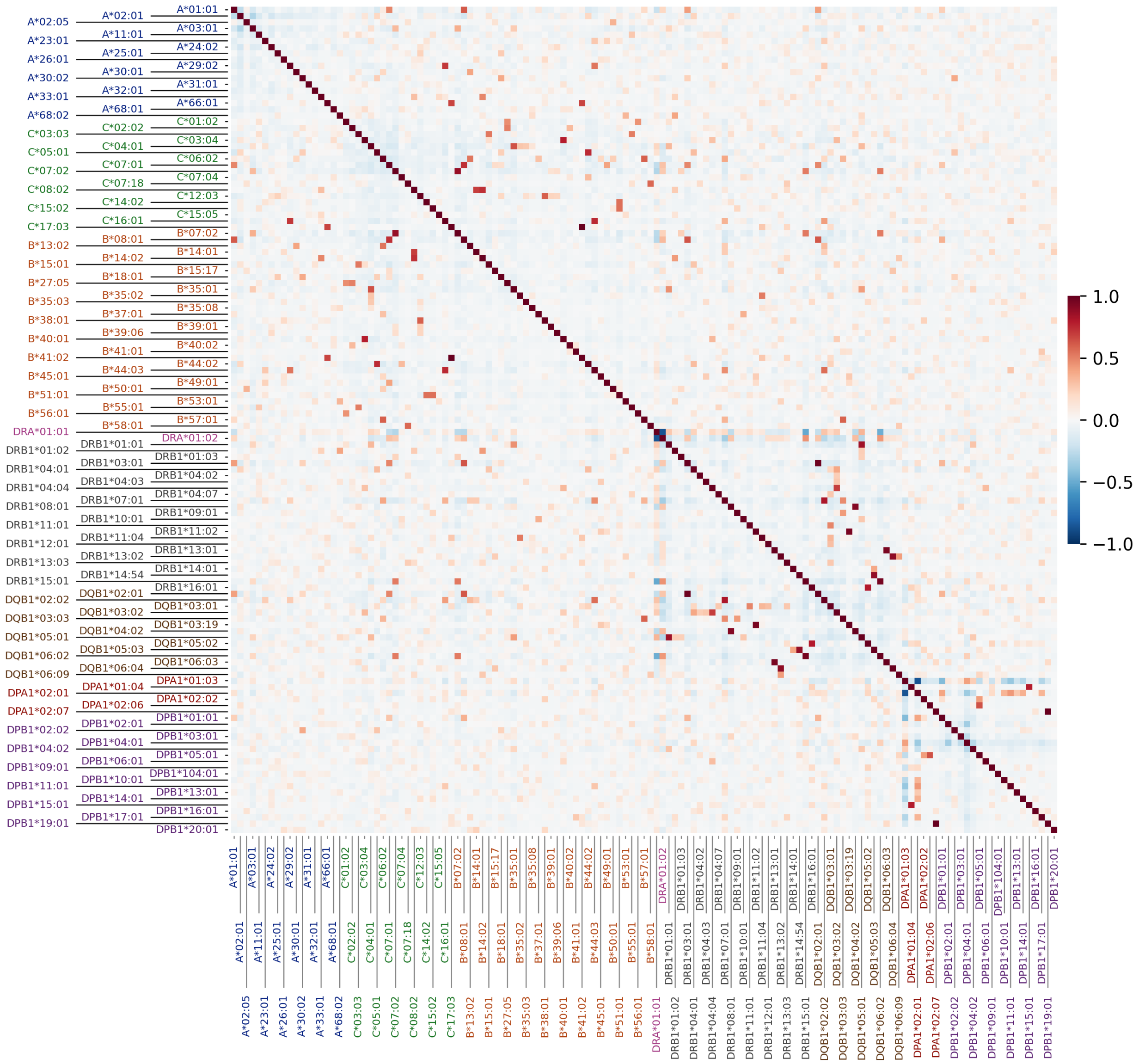


**Supplementary Figure 2D. Correlation between classical 2-field HLA alleles in the European genetic ancestry group.** Blue cells indicate negative Pearson correlation coefficients, red cells indicate positive Pearson correlation coefficients, and white cells indicate zero or near-zero correlation between allele pairs. Alleles are ordered alphanumerically by name. Alleles belonging to the same HLA gene are shown in the same color.

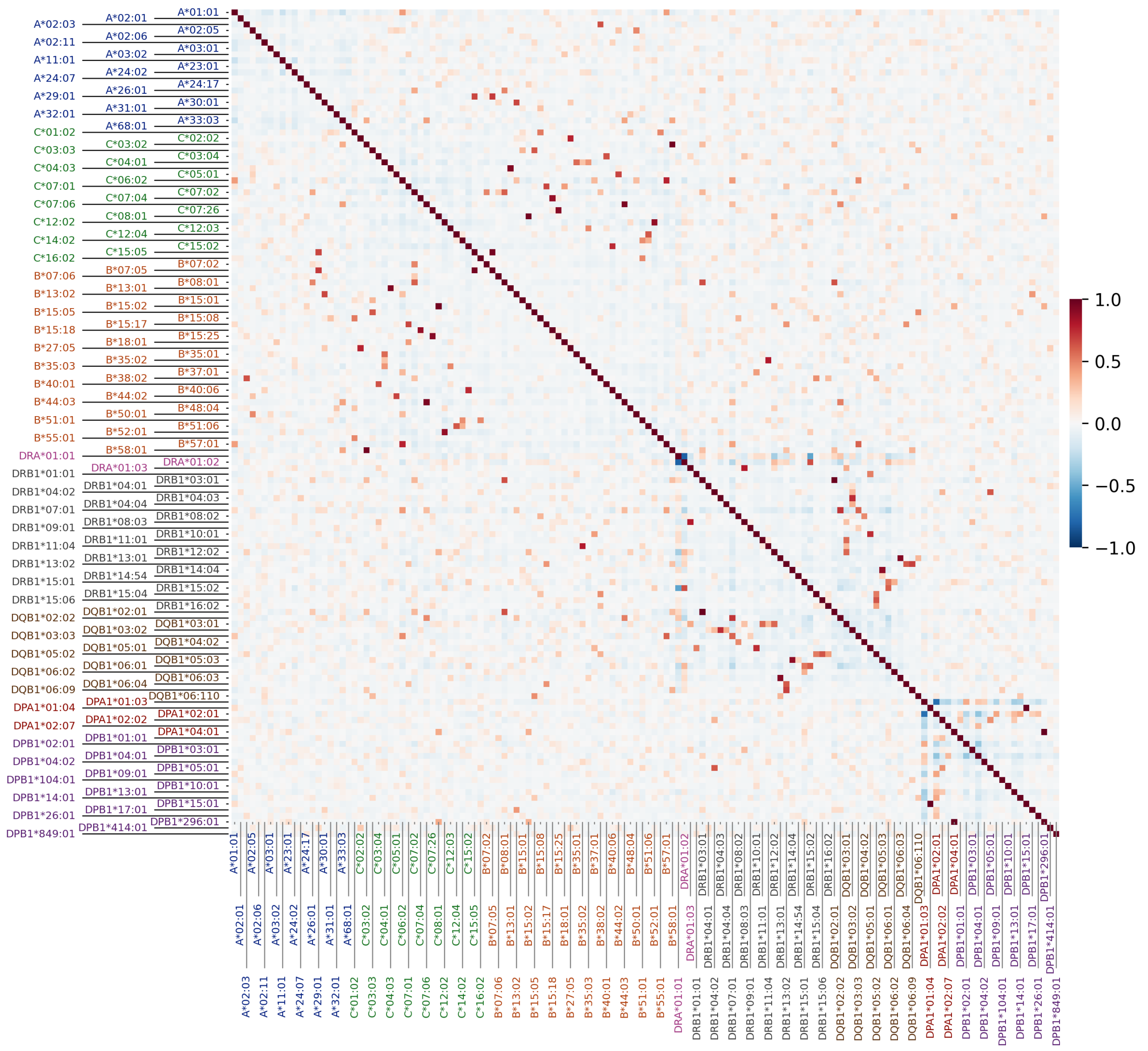


**Supplementary Figure 2E. Correlation between classical 2-field HLA alleles in the South Asian genetic ancestry group.** Blue cells indicate negative Pearson correlation coefficients, red cells indicate positive Pearson correlation coefficients, and white cells indicate zero or near-zero correlation between allele pairs. Alleles are ordered alphanumerically by name. Alleles belonging to the same HLA gene are shown in the same color.

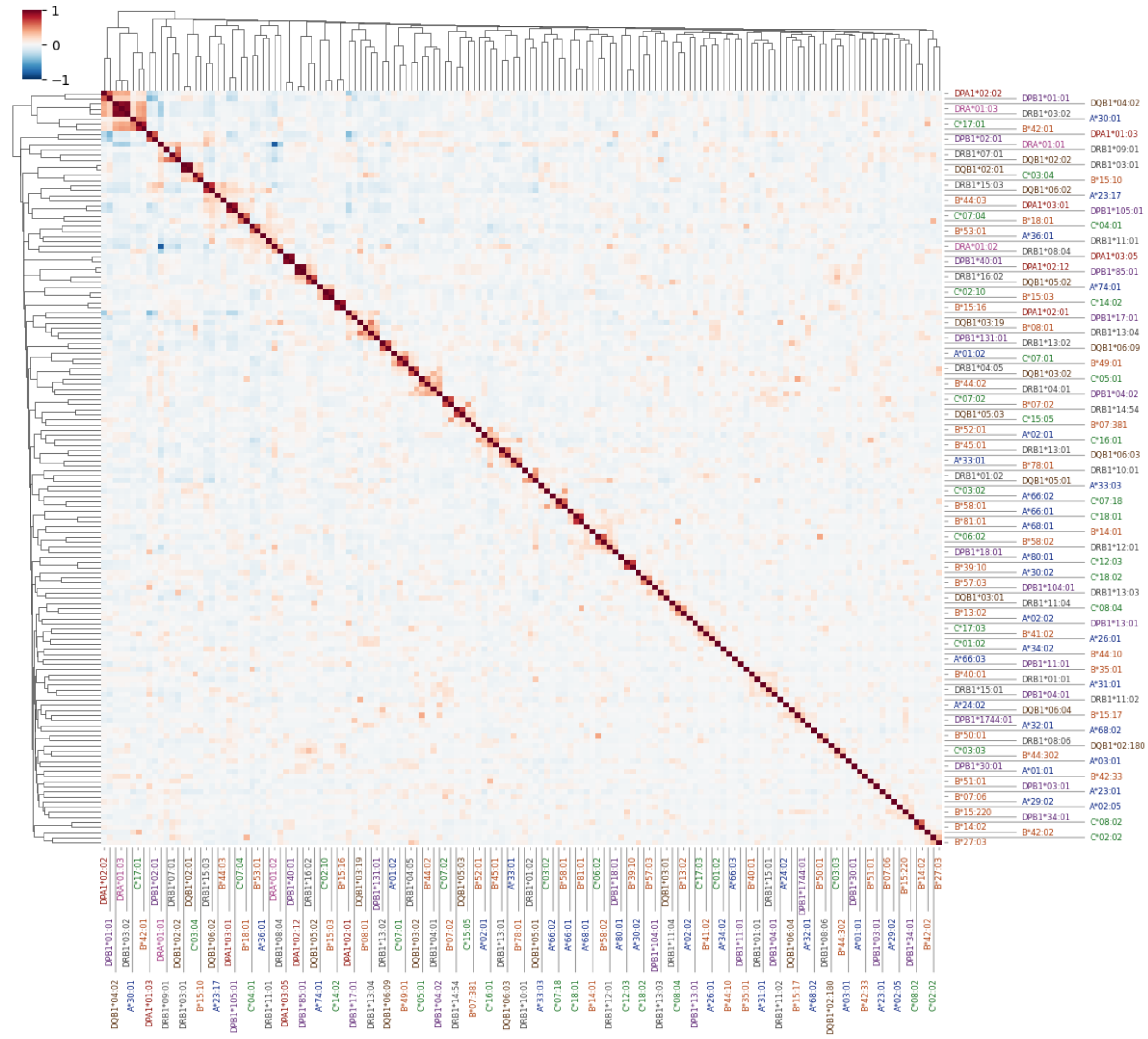


**Supplementary Figure 3A. Hierarchical clustering of classical 2-field HLA alleles in the African genetic ancestry group based on pairwise correlations.** Blue cells indicate negative Pearson correlation coefficients, red cells indicate positive Pearson correlation coefficients, and white cells indicate zero or near-zero correlation between allele pairs. Alleles are ordered according to hierarchical clustering. Alleles belonging to the same HLA gene are shown in the same color.

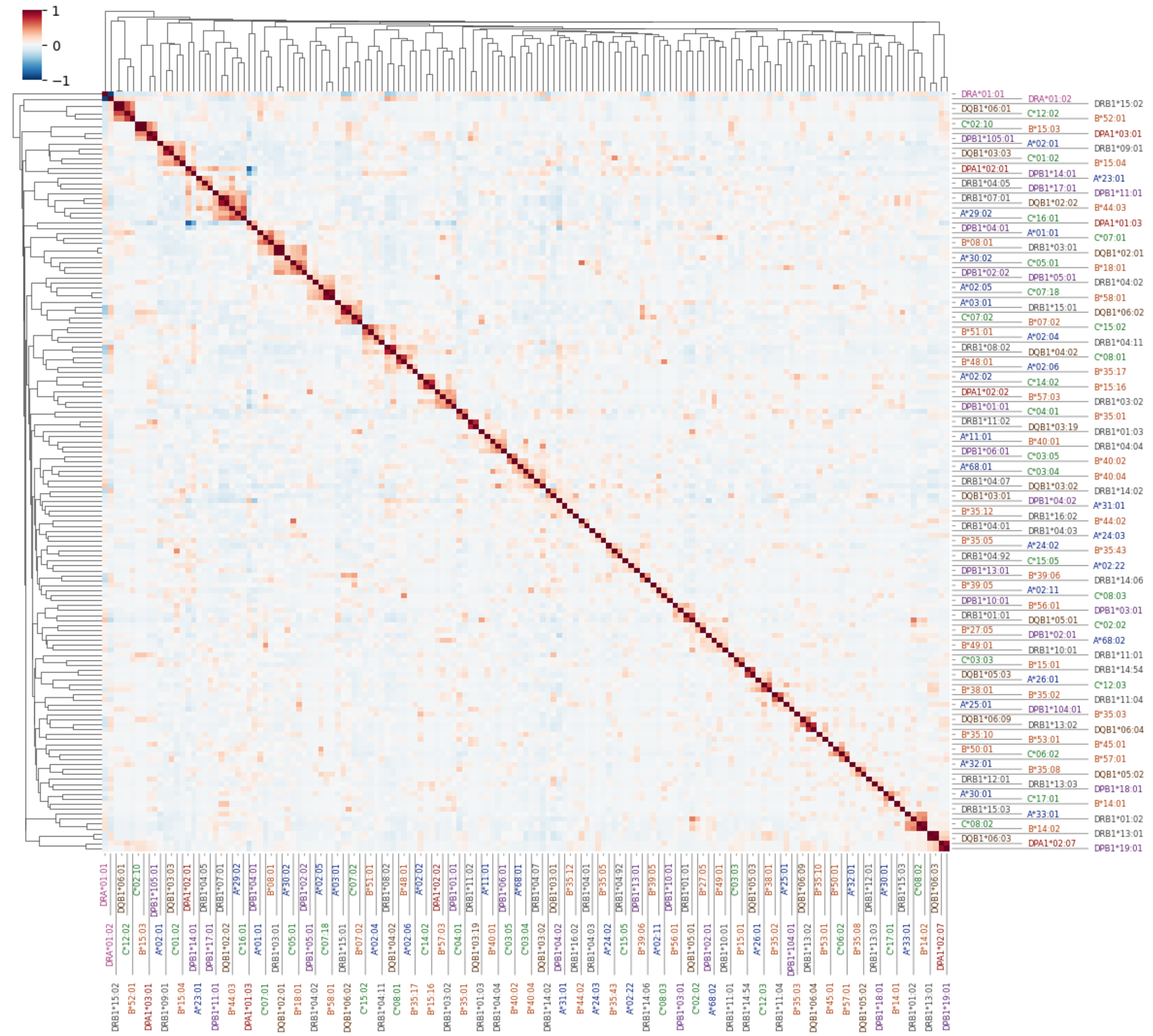


**Supplementary Figure 3B. Hierarchical clustering of classical 2-field HLA alleles in the Admixed American genetic ancestry group based on pairwise correlations.** Blue cells indicate negative Pearson correlation coefficients, red cells indicate positive Pearson correlation coefficients, and white cells indicate zero or near-zero correlation between allele pairs. Alleles are ordered according to hierarchical clustering. Alleles belonging to the same HLA gene are shown in the same color.

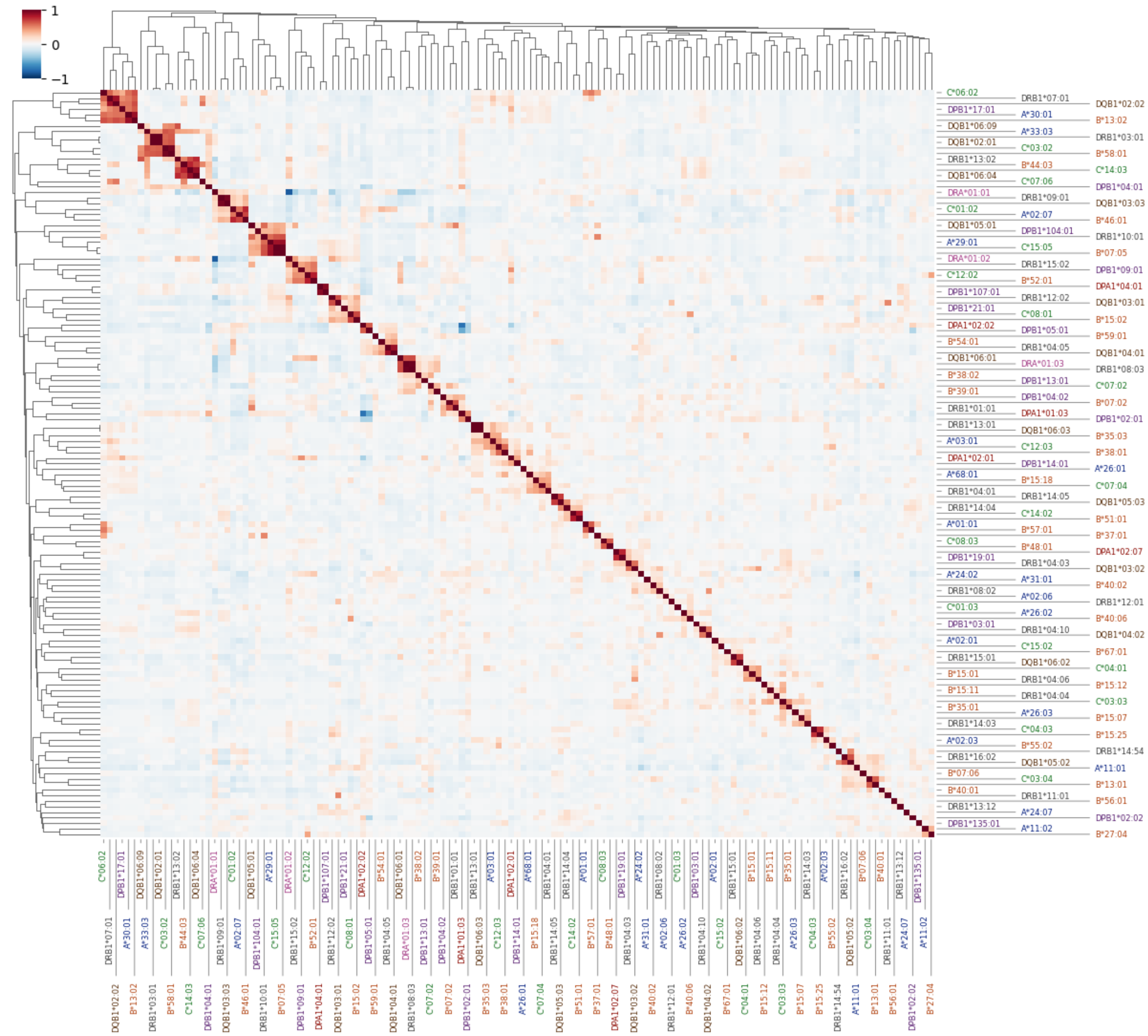


**Supplementary Figure 3C. Hierarchical clustering of classical 2-field HLA alleles in the East Asian genetic ancestry group based on pairwise correlations.** Blue cells indicate negative Pearson correlation coefficients, red cells indicate positive Pearson correlation coefficients, and white cells indicate zero or near-zero correlation between allele pairs. Alleles are ordered according to hierarchical clustering. Alleles belonging to the same HLA gene are shown in the same color.

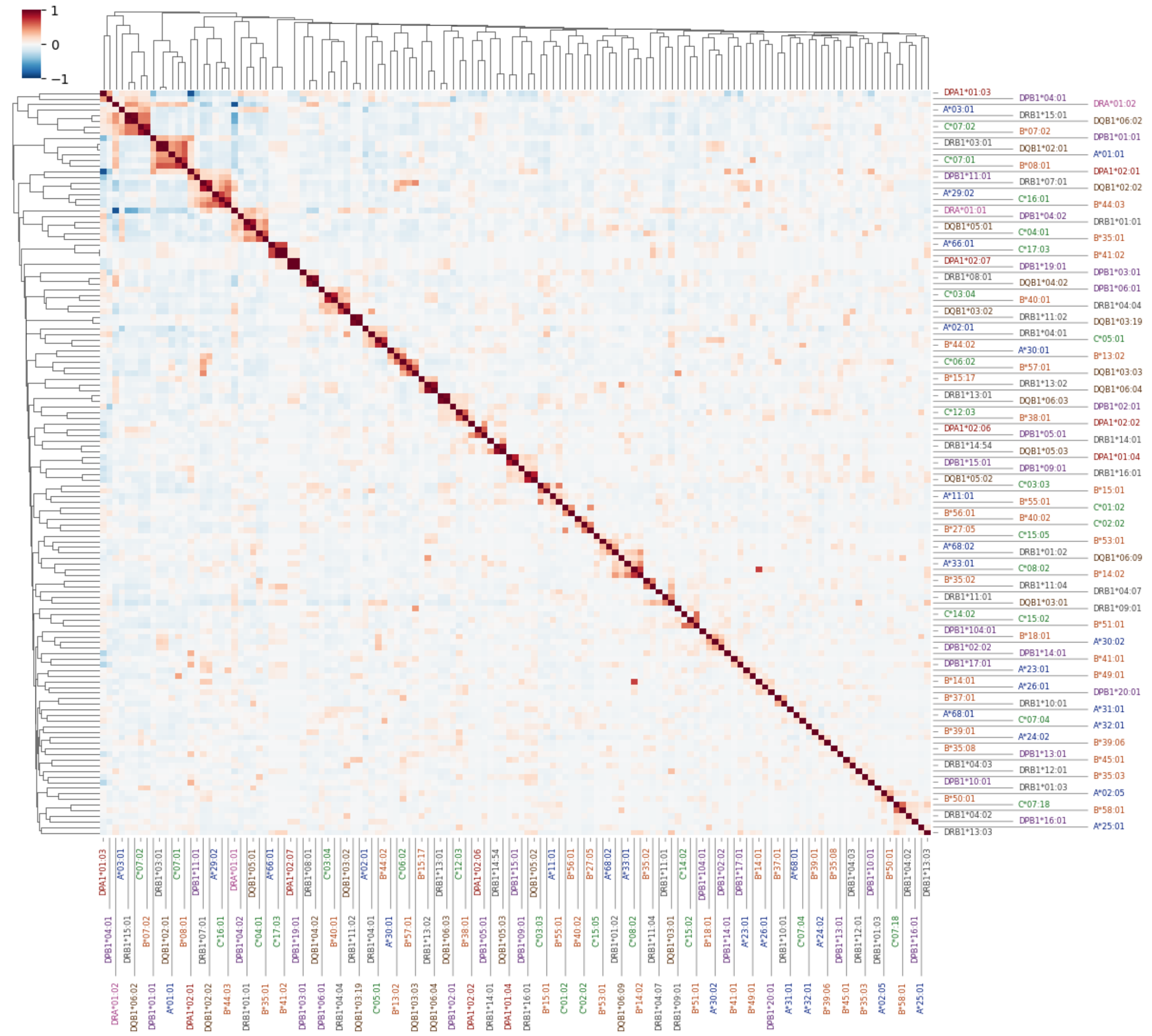


**Supplementary Figure 3D. Hierarchical clustering of classical 2-field HLA alleles in the European genetic ancestry group based on pairwise correlations.** Blue cells indicate negative Pearson correlation coefficients, red cells indicate positive Pearson correlation coefficients, and white cells indicate zero or near-zero correlation between allele pairs. Alleles are ordered according to hierarchical clustering. Alleles belonging to the same HLA gene are shown in the same color.

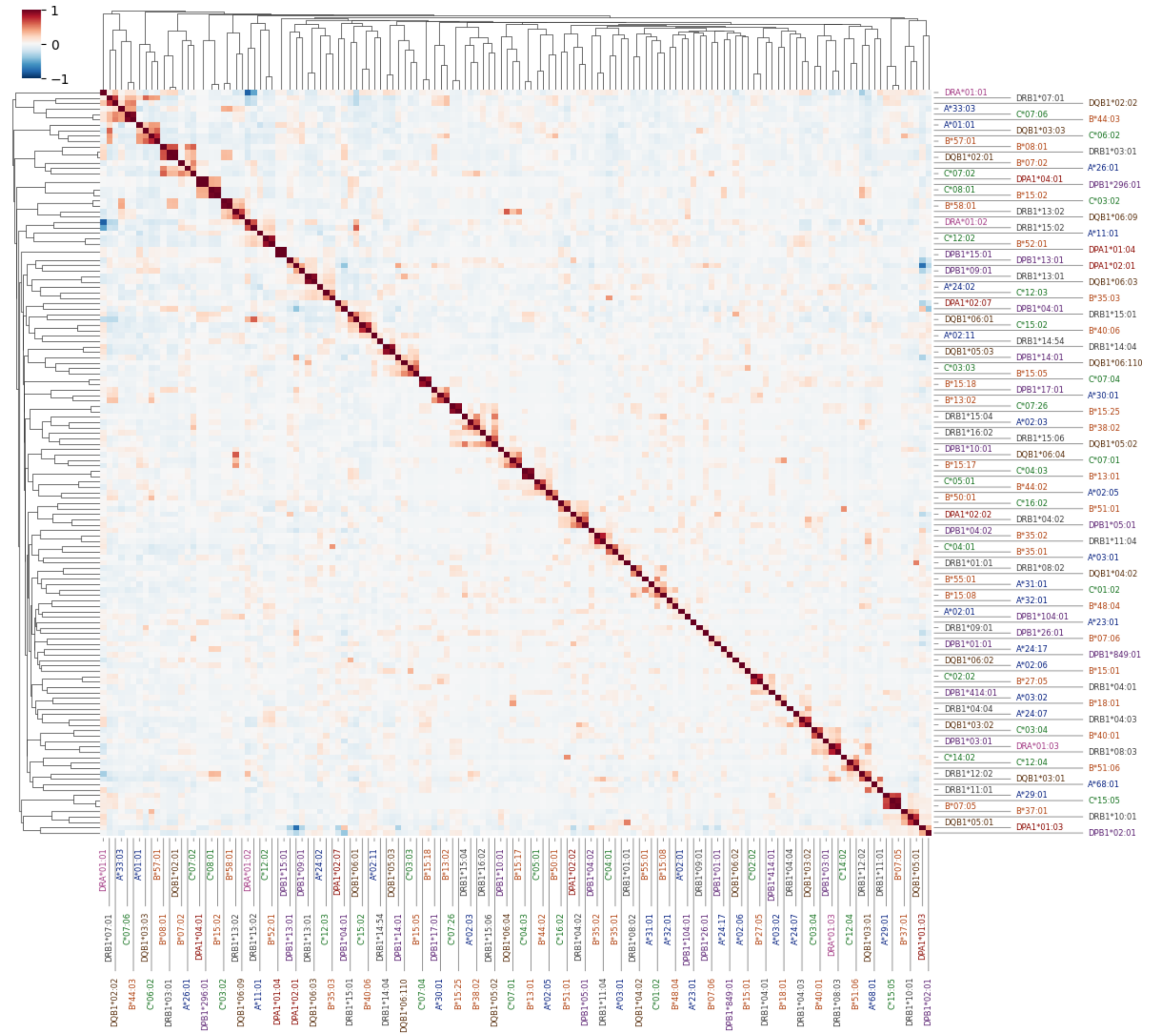


**Supplementary Figure 3E. Hierarchical clustering of classical 2-field HLA alleles in the South Asian genetic ancestry group based on pairwise correlations.** Blue cells indicate negative Pearson correlation coefficients, red cells indicate positive Pearson correlation coefficients, and white cells indicate zero or near-zero correlation between allele pairs. Alleles are ordered according to hierarchical clustering. Alleles belonging to the same HLA gene are shown in the same color.

**A) African genetic ancestry group**

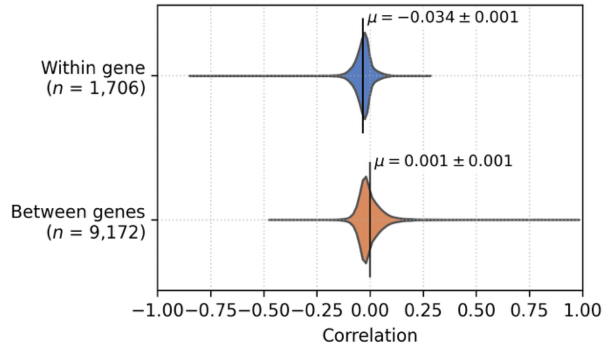


**B) Admixed American genetic ancestry group**

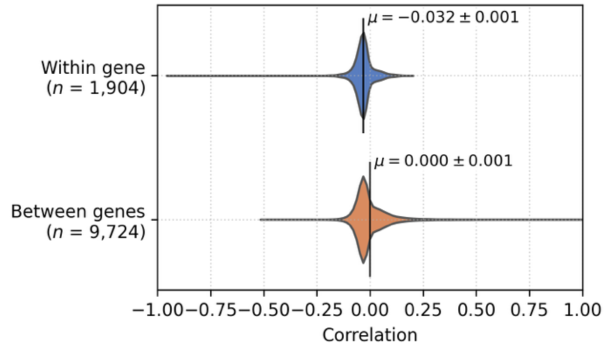


**C) East Asian genetic ancestry group**

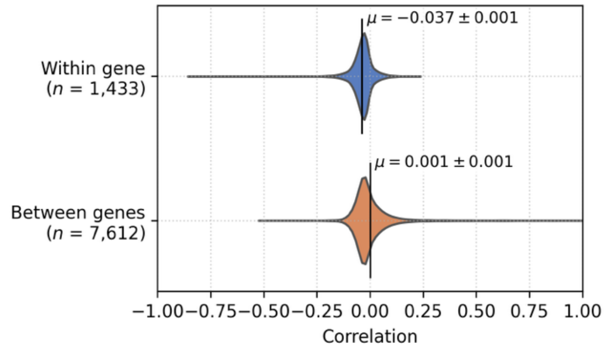


**D) European genetic ancestry group**

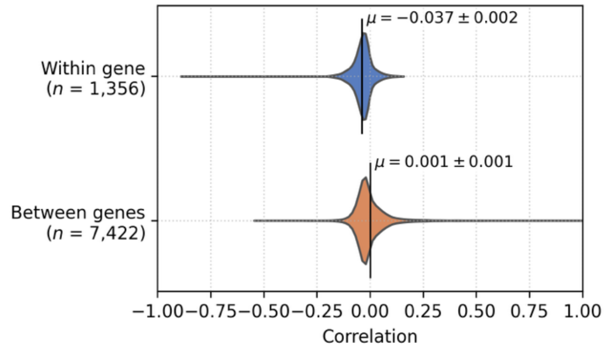


**E) South Asian genetic ancestry group**

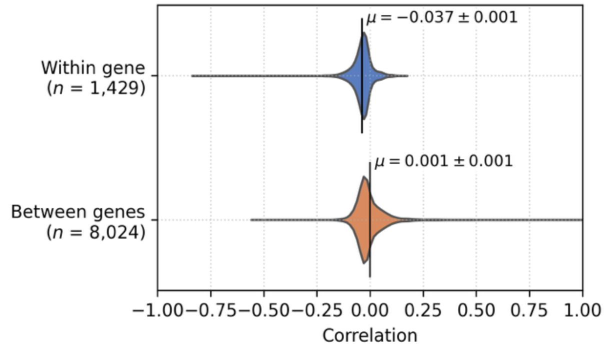


**Supplementary Figure 4. Distribution of pairwise Pearson correlation coefficients between 2-field HLA alleles within and between HLA genes for each genetic ancestry group.** Vertical lines indicate the mean correlation coefficient for each distribution, with annotated mean values and standard errors.

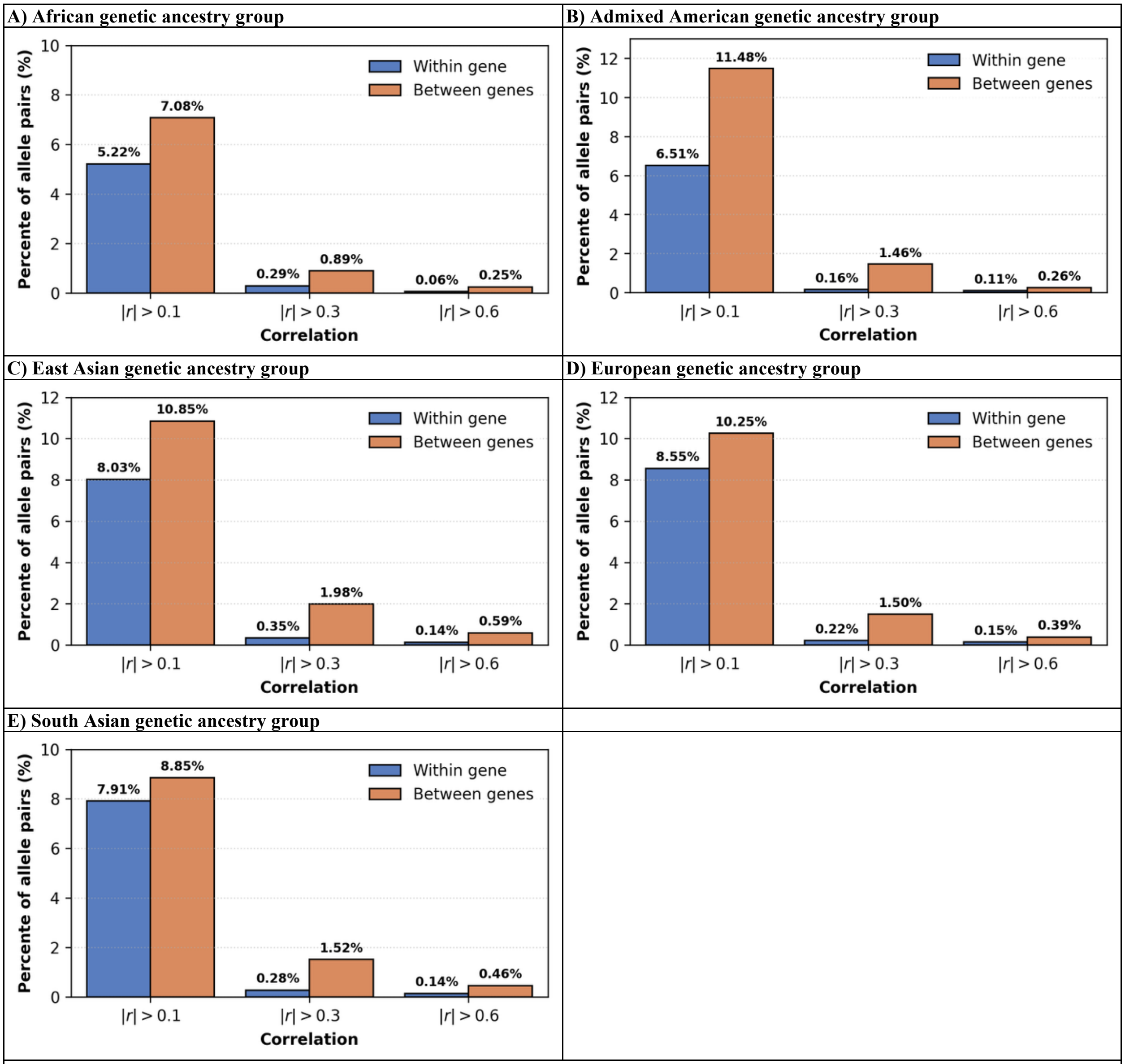


**Supplementary Figure 5. Percentage of 2-field HLA allele pairs with pairwise Pearson correlation coefficients exceeding selected thresholds within and between HLA genes for each genetic ancestry group.** Thresholds were applied to the absolute values of Pearson correlation coefficients (|r|). Bars are ordered by increasing correlation threshold.

**A) African genetic ancestry group**

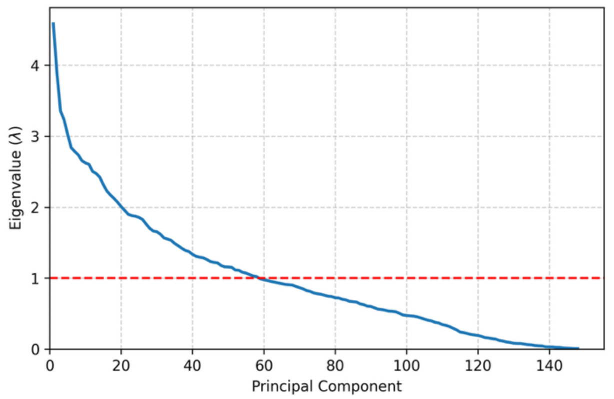


**B) Admixed American genetic ancestry group**

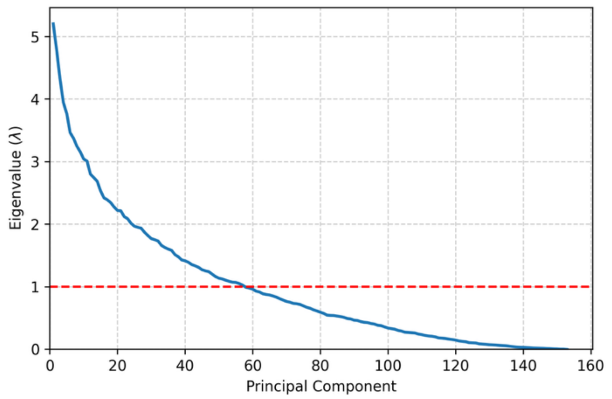


**C) East Asian genetic ancestry group**

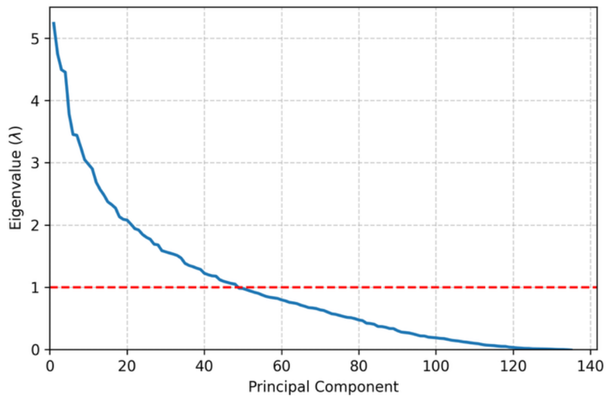


**D) European genetic ancestry group**

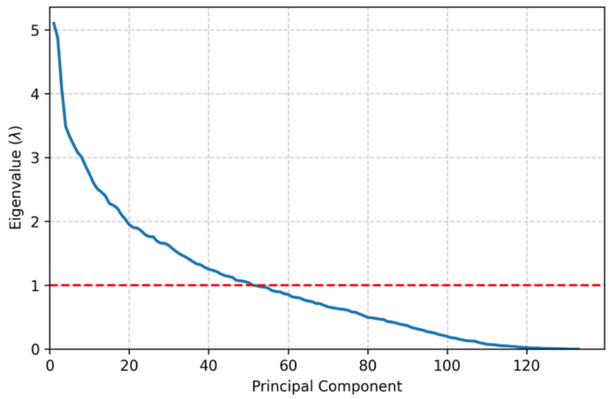


**E) South Asian genetic ancestry group**

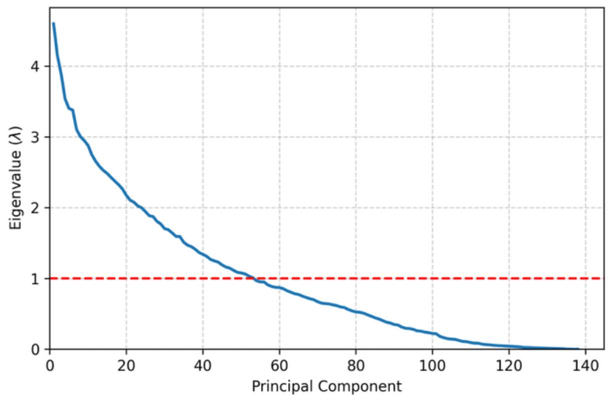


**F) All genetic ancestries combined**

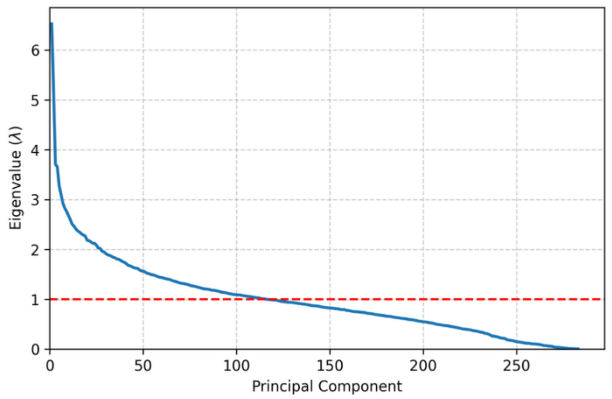


**Supplementary Figure 6. Scree plots (eigenvalue vs. principal component) for each genetic ancestry group.** Eigenvalue decomposition was performed on the pairwise Pearson correlation matrix of HLA alleles. Principal components are ordered by eigenvalue from largest to smallest. The red dashed line indicates the Kaiser criterion (λ = 1): principal components with eigenvalues greater than 1 explain more variance than an individual original variable.

**A) African genetic ancestry group**

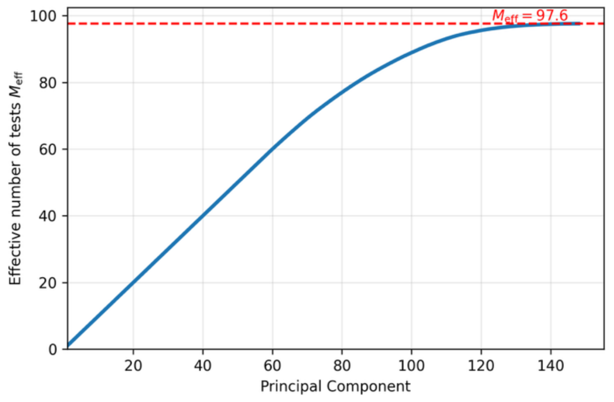


**B) Admixed American genetic ancestry group**

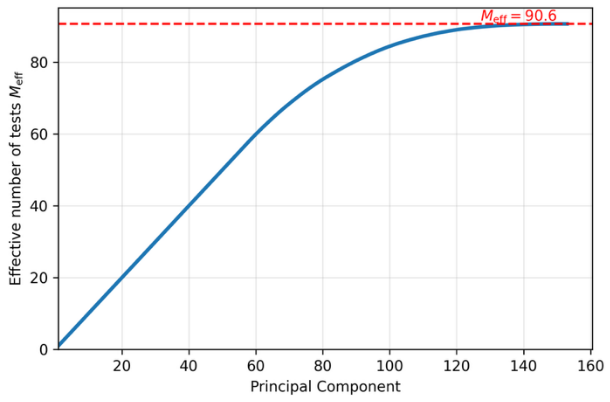


**C) East Asian genetic ancestry group**

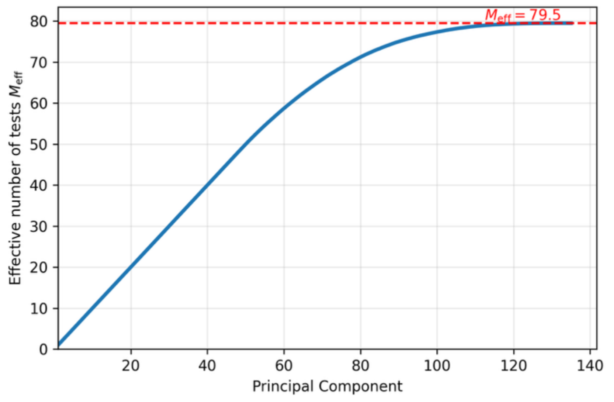


**D) European genetic ancestry group**

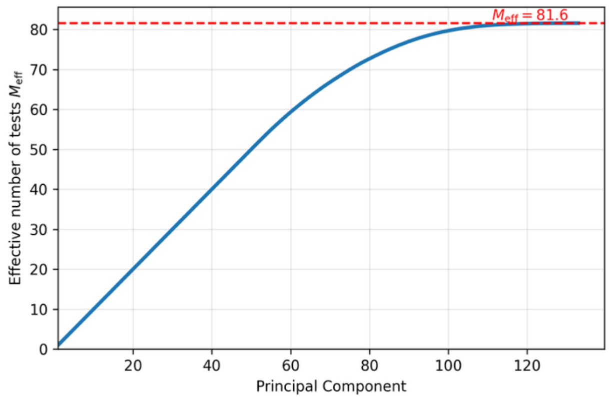


**E) South Asian genetic ancestry group**

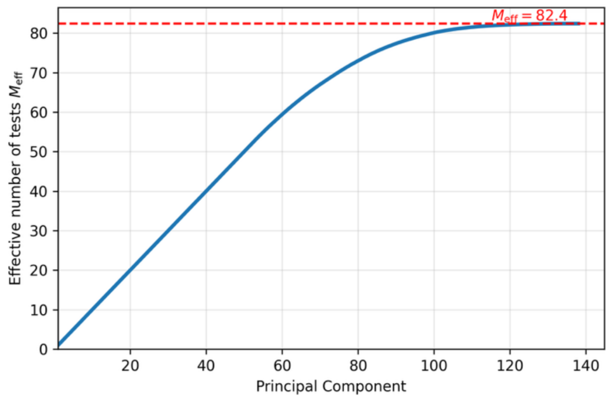


**F) All genetic ancestries combined**

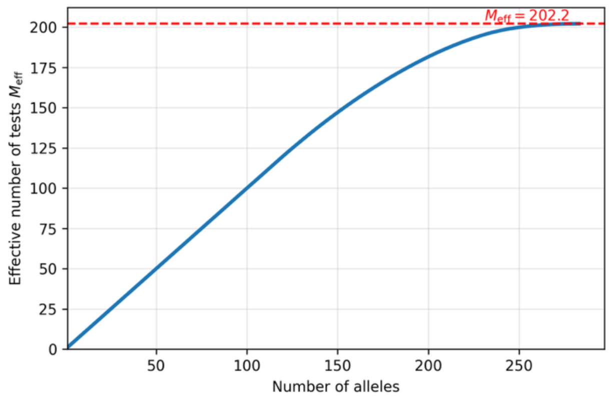


**Supplementary Figure 7. Eigenvalue-based effective number of tests as a function of principal components for each genetic ancestry group.** Eigenvalue decomposition was applied to the Pearson correlation matrix of HLA alleles, with components ordered by decreasing eigenvalue. Eigenvalue-based effective number of tests was estimated using the method of Li & Ji (2005). The red dashed line indicates the final eigenvalue-based effective number of tests.

| A. | B. |
|---|---|
| 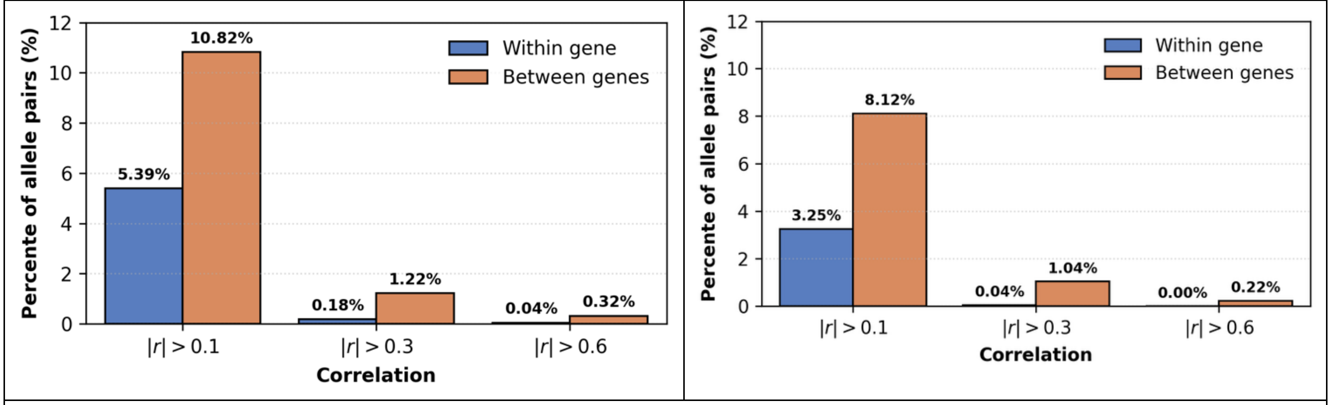 | |



**Supplementary Figure 8. Correlation between 2-field and 4-field HLA alleles typed in the 1000 Genomes Project's long read sequencing data.** Results are based on 415 individuals representing different genetic ancestries after quality control. **A.** 2-field HLA alleles. **B.** 4-field HLA alleles.
